# Distinct developmental mechanisms influence sexual dimorphisms in the milkweed bug *Oncopeltus fasciatus*

**DOI:** 10.1101/2021.05.12.443917

**Authors:** Josefine Just, Mara Laslo, Ye Jin Lee, Michael Yarnell, Zhuofan Zhang, David R. Angelini

**Affiliations:** Department of Biology, Colby College, 5700 Mayflower Hill, Waterville, ME 04901; Department of Organismic & Evolutionary Biology, Harvard University, 16 Divinity Avenue, Cambridge, MA 02138; Department of Stem Cell and Regenerative Biology, Harvard University, 7 Divinity Avenue, Cambridge, MA 02138; Department of Pediatrics, University of Colorado School of Medicine, 13123 E. 16th Ave., B065, Aurora, CO 80045; School of Electrical and Computer Engineering, Georgia Institute of Technology, 777 Atlantic Dr, Atlanta, GA 30332

**Keywords:** *Oncopeltus fasciatus*, sexual dimorphism, sex determination, *doublesex*, *intersex*, *fruitless*

## Abstract

Sexual dimorphism is common in animals. The most complete model of sex determination comes from *Drosophila melanogaster*, where the relative dosage of autosomes and X chromosomes leads indirectly to sex-specific transcripts of *doublesex*. Female Dsx interacts with a mediator complex protein encoded by *intersex* to activate female development. In males the transcription factor encoded by *fruitless* promotes male-specific behavior. The genetics of sex determination have been examined in a small number of other insects, yet several questions remain about the pleisomorphic state. Is *doublesex* required for female and male development? Is *fruitless* conserved in male behavior or morphology? Are other components such as *intersex* functionally conserved? To address these questions, we report expression and functional tests of *doublesex, intersex* and *fruitless* in the hemipteran *Oncopeltus fasciatus*, characterizing three sexual dimorphisms. *doublesex* prevents intersex phenotypes in all sexes and dimorphic traits in the milkweed bug. *intersex* and *fruitless* are expressed across the body, in females and males. *fruitless* and *intersex* also affect the genitalia of both sexes, but have effects limited to different dimorphic structures in different sexes. These results reveal roles for *intersex* and *fruitless* distinct from other insects, and demonstrate distinct development mechanisms in different sexually dimorphic structures.

## 1. Introduction

Sexual dimorphisms are wide-spread among animals, but their development has been studied in only a small number of species. These genetic model organisms have revealed a surprising degree of diversity in sex determination mechansisms [1–3], despite the deep evolutionary conservation of sex itself. However, sexual dimorphisms across the body are typically assumed to share similar regulation. Even in insects, where somatic sex determination is cell autonomous, similar mechanisms have traditionally been thought to control sex-specific differentiation in multiple tissues and organs [4–7]. Somatic sex determination requires an initial genetic signal, which varies from species to species [8–9]. Proteins downstream of this signal then regulate transcription sex-specifically.

The best-studied model of insect sex determination is *Drosophila melanogaster*, where a system of female and male chromosomal determinants reside on the X chromosome and autosomes, respectively [10–11]. Thus, the proportion of X chromosomes to autosomes indirectly determines sex. Sex-specific expression and splicing of downstream genes control female and male somatic sexual differentiation [12–14]. X-linked signal elements promote the expression of *Sex-lethal*, which directs female-specific splicing of *transformer* (*tra*) [15–16]. The female Tra protein is required for female-specific splicing of *doublesex* (*dsx*); without Tra a male-specific *dsx* transcript is produced [17–18]. The transcription factor isoforms encoded by *dsx* then regulate sex-specific development throughout the body [2,17,]. Female Dsx requires the mediator complex protein encoded by *intersex* (*ix*) as a cofactor [19–20]. Male-specific splicing is also required to produce the transcription factor encoded by *fruitless* (*fru*), which is necessary for male courtship behavior [14] as well as minor male anatomical features in *D. melanogaster* [21–23].

The evolution of gene regulatory networks is thought to be constrained by high degrees of epistasis and pleiotropy, which are expected to impose strong stabilizing selection [24]. Sex determination provides an intriguing exception to this pattern, with substantial variation even among Holometabola [1,3,25,26]. Amidst this variation, *dsx* is conserved in Holometabola as a point of integration for diverse sex-determining signals and *dsx* activity is required for the development of females and males [1,3,26–30]. However, outside Holometabola the functional conservation of *dsx* is unclear. RNA interference of *dsx* in the brown planthopper *Nilaparvata lugens* causes a pseudo-female phenotype in presumptive males, and minor effects in females who nevertheless retain normal fertility [31]. Knockdown experiments in the cockroach *Blattella germanica* [32] and in the brachiopod crustacean *Daphnia magna* [33] produced female traits in presumptive males, but had no effects on presumptive females. These results suggest that the ancestral role of *dsx* in insects is in directing male development. However, when this transition occurred during the insect radiation is unclear. There may be conserved roles for *fru* in male-specific behaviors, as evidenced by knockdown experiments in the cockroach [34] and silkmoth [35]. While *ix* is only required for somatic sexual dimorphism in female *D. melanogaster* [20], it is required during genitalia development in females and males of the hemipteran *Oncopeltus fasciatus* [36], suggesting a broader role in the development of some insect groups for genes with more circumscribed functions in *D. melanogaster*.

Our understanding of the extent of conservation or divergence in insect sex determination is currently limited by the small number of species for which functional genetic tests have been undertaken. Here, we report the study of somatic sex determination in the milkweed bug *Oncopeltus fasciatus* (Heteroptera : Lygaeidae), focusing on *dsx* and two sex determination genes with divergent functions: *ix* and *fru*. The true bugs are a diverse and species-rich hemimetabolous order that diverged from holometabolous insects more than 370 million years ago [37], making them an important point of phylogenetic comparison for more well-studied species. Milkweed bug females and males differ in the structure of the genitalia, in the curvature of the fourth abdominal sternite, and in the distribution of abdominal melanization (figure 1). We find a common requirement for *dsx* activity in *O. fasciatus* females and males, in multiple sexually dimorphic structures. We find an unexpected, major role for *fru* in development of both female and male sexually dimorphic structures. Different dimorphic traits require distinct combinations of these genes. These findings indicate that the sex determination pathway of milkweed bugs has diverged significantly from that of other insects examined thus far.

**Figure 1.**
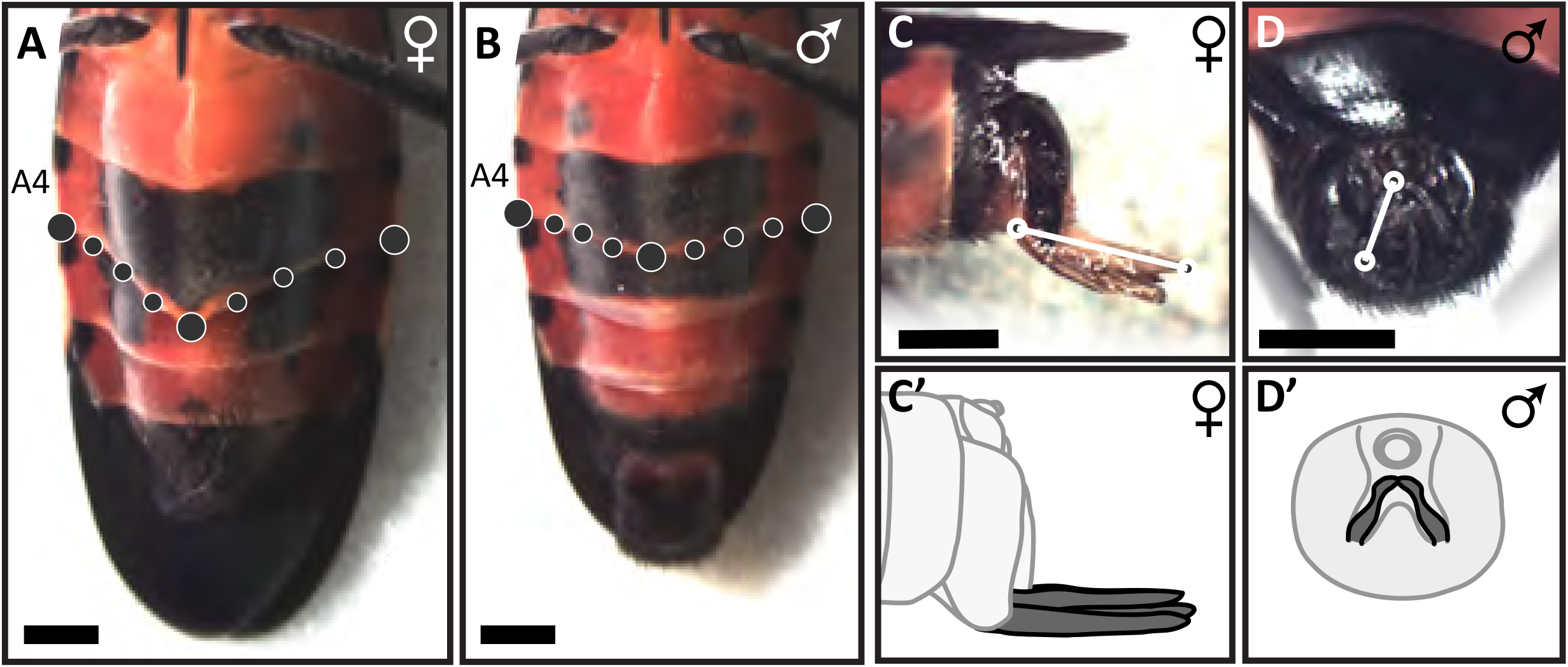
Milkweed bugs have several sexually dimorphic features. (a-b) The fourth abdominal body segment (A4) of females has an exaggerated extension of the sternite at the ventral midline. Nine landmarks were placed along the ventral edge of the sternite to quantify curvature. (c) Appendages from the terminal segments of the female abdomen form an ovipositor. Ovipositor length was measured from the distal tip to the proximal joint. (d) The male abdomen ends in a large genital capsule, which contains the copulatory organ. Externally a pair of small claspers, visible from a posterior perspective, is used for mate guarding. Clasper length was measured from the proximal joint to the tip.

## 2. Materials and Methods

### (a) Insect culture

*Oncopeltus fasciatus* were obtained from Carolina Biological Supply Company and maintained as described by Liu and Kaufman [38]. Milkweed bug cultures were kept in terraria at room temperature (21-23°C) and fed on sunflower seeds. Loose cotton balls provided an egg-laying substrate. Water was provided in flasks with paper towel wicks.

### (b) Sequence orthology

An ortholog of *ix* from *O. fasciatus*, GenBank accession AEM16993, was identified as part of a previous study [36]. Sequences with similarity to *fru* and *dsx* were obtained from *O. fasciatus* using degenerate PCR and RACE and by PCR with exact primers based on the available genome and gene set annotation [39–40]. Initial identification was based on tBLASTn search using sequences from *D. melanogaster*. One genomic locus had high similarity to *fru*, while three genomic loci closely resembled *dsx*.

Orthology was confirmed using phylogenetic inference (figures S1-S4). Additional orthologs from insects were obtained from GenBank using BLAST. Alignment of amino acid sequences was performed by ClustalOmega version 1.2.4 [41] using two iterations. Alignment included all amino acids. Phylogenies were inferred using RAxML version 8.2.11 [42]. Mutation rates were modeled using the BLOSUM62 matrix with a gamma distribution. Node supports were determined by bootstrapping. The program was called as “`raxml-mpi -d -f a -x 123 -p 123 -# autoMRE -m PROTCATBLOSUM62 -s X.phy -n X`” where `X` stands in for the alignment name.

### (c) Isolation of mRNA isoforms

Alternative splicing of mRNAs was examined using RACE (rapid amplification of cDNA ends). Initial cloning of candidate genes was done via degenerate PCR with primers designed to conserved protein domains, or with exact primers designed from publicly available transcriptome data. To obtain alternative mRNA sequences, total RNA was extracted from a mix of late fifth instars and teneral adults seperated by sex, using the PureLink RNA Mini Kit (ThermoFisher). First strand cDNA synthesis was done using the GeneRacer Kit with AMV reverse transcriptase (Invitrogen). This procedure ligates a specific DNA adapter sequence to the 5′ or 3′ ends of cDNAs, which are then used in PCR with gene-specific and adapter-specific primers. Nested PCR was used to enrich the template pool for the target sequences. From multiple independent PCRs, 120 products were isolated using gel excision and cloned using the Topo-TA cloning kit (Invitrogen). Sanger sequencing and alignment confirmed 51 of these clones had contiguity with DMRT genes. These were determined to represent 3 genomic loci with a total of 11 splicing isoforms (figure 2a-b; S6a-b). 5′ RACE using gene-specific reverse primers in more downstream exons failed to isolate additional transcript variants. BLAST searches of available *O. fasciatus* transcriptomes (Ewen-Campen et al. 2011; Panfilio et al. 2019) using individual exons as a query also failed to identify additional transcripts. Sequences from *O. fasciatus* transcripts were submitted to GenBank (accession numbers MZ197788 - MZ197800).

**Figure 2.**
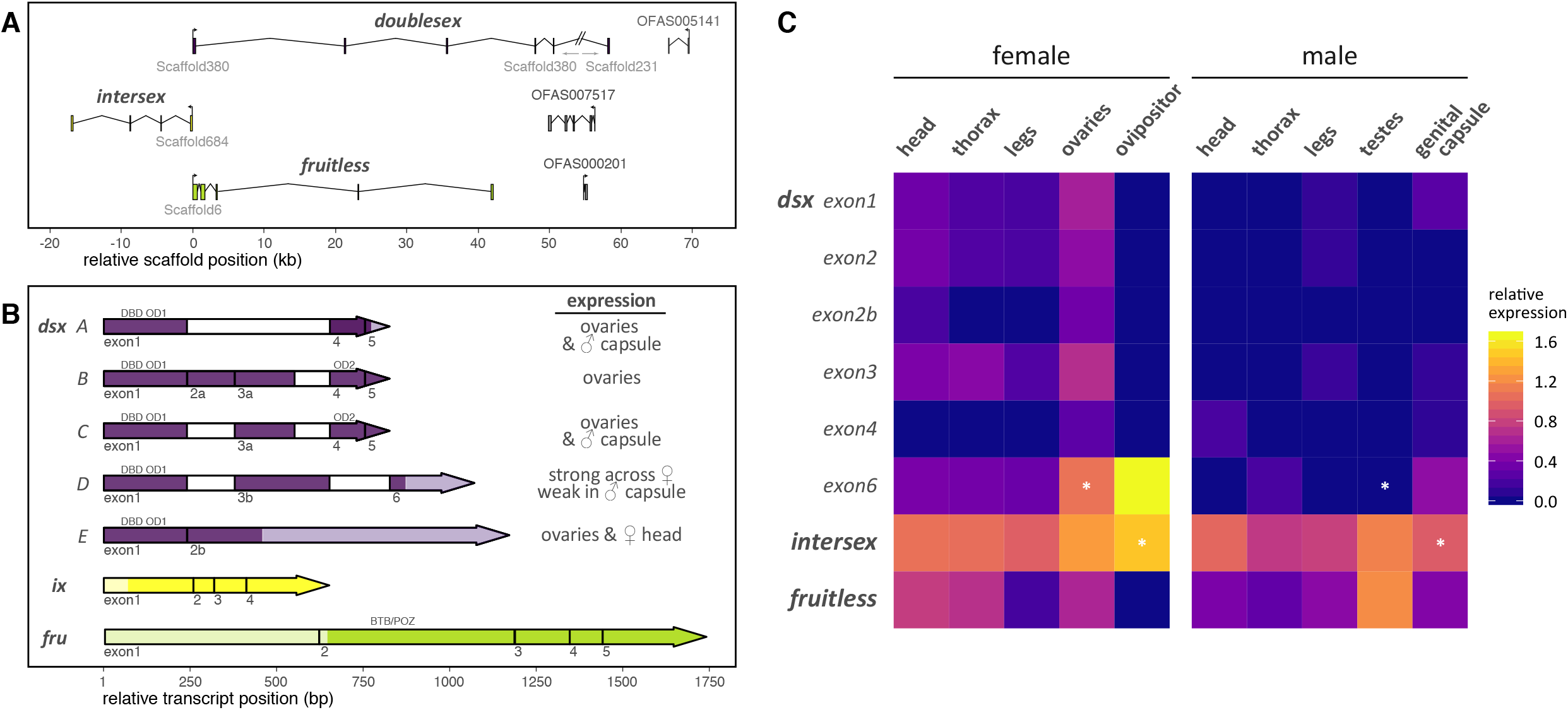
Structure and expression of *doublesex, intersex* and *fruitless* in *O. fasciatus*. (a) Orthologs were identified on different genomic scaffolds in the *O. fasciatus* genome. Boxes represent exons; angled lines represent introns. Introns were very long compared to neighboring gene predictions. Flags indicate the start of transcription. (b) Transcripts from *doublesex, intersex* and *fruitless*. The first exon of each transcript is aligned at the left. Omitted exons are left empty. Alternative start or stop sites for exons are indicated by lowercase letters (e.g. 2a, 2b) Lighter shading indicates UTRs. Conserved protein domains are noted above the exon. (c) Expression of candidate sex determination genes in *O. fasciatus*. A heatmap shows the relative transcript abundance from five body regions in each sex for *doublesex*, *intersex* and *fruitless*. Each panel represents the mean of 5 to 7 independent measurements. Expression differences between females and males were tested using the Wilcoxon rank sum test (* indicates *p* < 0.05).

### (d) Gene expression

Quantitative real-time PCR (qRT-PCR) was used to measure sex- and trait-specific expression, to validate RNAi knockdowns, and to investigate potential gene interactions. Specific probe and primer sets were created for multiple exons of *doublesex*. Dual-labeled probes (SigmaAldrich) allowed reaction multiplexing to conserve template samples. Primers and probes were designed using Primer3 in Geneious Prime and are listed in Supplemental Table 1.

Milkweed bugs were collected without agitation and placed at −80 °C for fifteen minutes. For tissue-specific assays, specimens were dissected on ice and immediately proceeded into RNA isolation. RNA was extracted following the Maxwell 16 LEV Tissue RNA Kit protocol (Promega). For tissue-specific assays, RNAi validation and studies of gene interactions, expression was measured in templates prepared from teneral adults. Between five and seven biological replicates of each sex-by-tissue combination were measured independently. RNA was stored at −80 °C. First-strand cDNA was generated from 1 μg total RNA using the iScript cDNA kit (BioRad) with a poly-T primer as described in the kit’s instructions. qRT-PCR was carried out on a CFX96 Touch qPCR System (BioRad) using iQ Supermix (BioRad). Expression was calculated from the mean of technical triplicate reactions. Each plate included four DNA standards diluted from known concentrations of synthesized gene fragments (IDT gBlocks) with the addition of nonspecific salmon sperm DNA (ThermoFisher). All plates included control reactions with water in place of the cDNA template (no-template controls) and with total RNA in place of cDNA (no-RT controls). Whenever primer sets were used on multiple plates, a minimum of two templates (in addition to concentration standards) were included on those plates to allow inter-plate calibration. *Elongation Factor 1-α* (*EF1-α*) was used as an endogenous reference gene, chosen for its relative consistency throughout development [43].

We expected that in some tissues, certain transcripts would not be expressed. Therefore, we developed a method for calculating normalized gene expression that allows zero values to be determined in comparison to negative controls from the same plate. Typically, normalized target gene expression is determined by subtracting the mean C_q_ for a reference gene from the mean C_q_ of the gene of interest. In such cases, zero has no special meaning. Instead, we coded all measurements of gene expression as zero if they were within 0.5 cycles of any template-negative controls on the same plate, and maintained that value throughout analyses. Non-zero values were normalized as usual to the expression of *EF1-α*. Next, expression values were scaled, so that the mean of non-zero values was standardized to 1. Finally, variance was scaled to maintain the relative distance from zero of the global non-zero minimum value prior to standardization.

### (e) RNA interference

Loss-of-function phenotypes from RNA interference (RNAi) were used to investigate gene function as previously described [36]. Knockdown was verified by qRT-PCR. Double-stranded RNA (dsRNA) encoding *Ampicillin Resistance* (*AmpR*) was used as a control for dsRNA toxicity and injection mortality. This sequence was chosen because it is present on the pCR-Topo4 plasmid vector used in cloning of RACE gene fragments.

A synthetic gene fragment (IDT gBlock) or cloned gene fragment was used to create a PCR template for the synthesis of dsRNA. PCR primers were designed with an approximately 20-bp exact match to the target sequence, avoiding sequences that also occur in paralogs (table S2). A 5′ T7 polymerase promoter sequence was added to both primers. Following PCR, dsRNA was transcribed using the T7 MegaScript Transcription Kit (ThermoFisher). Template DNA was removed by treatment with DNase I. Slow cooling to room temperature allowed dsRNA annealing. Samples were cleaned by precipitation in cold ethanol and ammonium acetate. The product was resuspended in nuclease-free water and stored at −20 °C. Double-stranded RNA structure was confirmed by a coherent band on an agarose electrophoretic gel. dsRNA concentrations were measured using a microvolume spectrophotometer. dsRNA solutions were diluted in saline buffer (0.01 mM NaPO_4_, 5 mM KCl, green food coloring) and stored at −20 °C.

RNAi was tested at several dsRNA concentrations (table S3). Milkweed bugs have five juvenile instars, and fourth instar *O. fasciatus* are not yet sexually dimorphic. Fourth instars were anesthetized using a CO_2_ pad and injected with dsRNA solutions using a borosilicate glass microcapillary, pulled on a Sutter Instruments pipette puller into an injection needle. Approximately 0.5 μl of dsRNA solution was injected into the left side of the abdomen of each bug.

The effectiveness of RNAi was validated using qRT-PCR. Gene expression was measured in control (*AmpR* dsRNA) specimens and RNAi treatments targeting the genes of interest (figure S5) in adults no more than 1 day old. Primers used for validation PCRs did not overlap with the dsRNA sequences. RNAi validation of *dmrt99B* RNAi was not possible, because expression of *dmrt99B* could not be detected in the whole bodies of 28 *AmpR* or 16 *dmrt99B* RNAi specimens examined. While *dmrt99B* expression was detected in specific body regions of unmanipulated bugs (figure S6c), the overall expression of *dmrt99B* appears to be very low.

### (f) Morphometry

We examined three somatic sexual dimorphisms, genitalia length, sternite curvature, and abdominal pigmentation patterns (figure 1). Specimens were imaged on a VWR VistaVision dissecting microscope with a Moticam 5 digital camera. A ruler was included in the image for scale. The ImageJ [44] line-tool was used to measure the length of genitalia and the femur, which was used to normalize for body size. In females, the absolute length of the first valvulae of the ovipositor was measured from the point of emergence to the distal tip (figure 1c). For males, the segmented line-tool was used to measure clasper length from the proximal joint to the tip (figure 1d). Using the multi-point tool, nine points were placed across the posterior edge of the fourth abdominal sternite (figure 1a-b). Raw pixel values were converted to metric distances based on measurement of the ruler scale in each image.

To quantify sternite curvature, landmark coordinate positions from each specimen were marked in ImageJ and stored in the TPS format [45]. Coordinates were then aligned using generalized Procrustes alignment, as implemented in the R package geomorph [46]. Landmarks at the left and right sides and on the anatomical midline were fixed, but others were treated as semilandmarks. Curvature was defined as the residual mean standard deviation (RMSD) from a linear regression of Procrustes-aligned coordinate values. Larger RMSD values indicated a higher amount of curvature in the sternite, the more characteristically female phenotype.

Ventral abdominal pigmentation of *O. fasciatus* adults varies with sex (figure 1a-b; 5g-i). Rearing temperature can also influence pigmentation [47]. Bugs in this study were kept at a consistent temperature (21 - 23°C), limiting this environmental influence. Within 1-2 hours of the imaginal molt, melanic areas are visible on the ventral abdomen. These areas were quantified using the freehand selection tool in ImageJ. Pixel areas were converted to area based on measurement of a mm scale in each photograph.

We examined the gonads of RNAi specimens but found no obvious phenotypic differences at the anatomical level (figure S7).

### (g) Statistical analyses

Measures of gene expression and quantitative phenotypes produced through RNAi are expected to follow skewed disruptions. Therefore, we used nonparametric statistical methods to analyze these data. We used the Kruskal-Wallis analysis of variance to detect overall effects. Individual contrasts were examined *post-hoc* using the two-sample Wilcoxon rank sum test. We used one-sided tests for the comparison of sex-specific phenotypes. Where appropriate, correction for multiple tests was done using the FDR method. R version 4.0.3 was used for all statistical analyses and the `ggplot2` package was used for plotting data. Figures were assembled in Adobe Illustrator.

## 3. Results

### (a) Identification of candidate sex determination genes in the milkweed bug

To examine the determination of somatic sexual dimorphisms in *O. fasciatus*, we cloned several candidate genes based on developmental models from *D. melanogaster* [4], including *dsx*, *ix* and *fru. dsx* is a key regulator of sex determination in many holometabolous insects [1,3,20,30]. *ix* is required for genitalia development in both sexes in *O. fasciatus* [36] but only in females of *D. melanogaster* [20]. In *D. melanogaster*, the transcription factor encoded by *fruitless* (*fru*) is required for sexually dimorphic brain development, necessary for male courtship behavior [14,21,48].

*doublesex* is a member of the *doublesex/mab-3* related transcription factor (DMRT) gene family [49], which in *D. melanogaster* includes *dmrt11E, dmrt93B* and *dmrt99B*, in addition to *dsx*. Three genomic loci in *O. fasicatus* have sequence similarity to *D. melanogaster* DMRT genes. We used phylogenetic inference to determine the orthology of these genes and to explore the relationships among them (figure S1). This analysis included 413 sequences from diverse insect lineages and non-insect DMRT outgroups. For each species included in the analysis, predicted protein sequences with strong BLAST similarity or annotations to any DMRT gene were included in the analysis. The maximum likelihood best tree (figure S1) forms four clades with modest bootstrap supports (45 - 69%) and excludes most non-insect DMRT sequences. Within each DMRT paralog clade the tree topologies do not recapitulate most well-accepted taxonomic relationships [26]. Disagreement with organismal relationships is also found for a phylogeny inferred from alignment of Dsx amino acid sequences alone (figure S2). In both trees, sequences from the same insect order are mostly grouped together, but node supports are not strong above the ordinal level.

Sequences from *O. fasciatus* are found in the *dsx*, *dmrt93B* and *dmrt99B* clades, closely associated with other sequences from Heteroptera (figure S1). No ortholog of *dmrt11E* was identified from *O. fasciatus*. Each of these proteins has an N-terminal region with a DNA-binding domain and a putative oligomerization domain (OD1) [50]. Each protein also has a conserved C-terminal region containing a second putative oligomerization domain (OD2). Due to the sequence similarities among DMRT genes, we performed parallel experiments using each *O. fasciatus* DMRT gene in case any had a role in sex determination in this species. We focus on results from *dsx* here; results from *dmrt99B* and *dmrt93B* did not reveal functions in sex determination and are therefore reported in the Supplemental Materials.

Single orthologs of *ix* and *fru* were found in most available insect genomes. Single copies of *ix* and *fru* were isolated from *O. fasciatus* (figure 2a). Transcriptome and genome sequences also supported single instances of these genes in the milkweed bug. The phylogeny inferred for *fru* amino acid alignment also deviates from the accepted relationships among major taxa (figure S3), however the *ix* phylogeny is closer to the expected relationships (figure S4) [26].

### (b) Structure- and sex-specific transcript expression

To effect sexually dimorphic development, genes are often alternatively spliced. Using rapid amplification of cDNA ends (RACE) we isolated transcripts for *dsx*, *fru*, and *ix* (figure 2b). We supplemented this survey by searching available transcriptome data, but this approach did not identify any additional transcripts. Five alternative transcripts were found for *O. fasciatus dsx*. All transcript isoforms from each *doublesex* paralog included a DNA-binding domain in the 5′ exon. An OD2 domain was present in the downstream exons of two transcript isoforms. We also identified three isoforms each from *dmrt93B* and *dmrt99B* (figure S6). Only a single transcript was identified for *ix* and *fru*. In *D. melanogaster fru* is spliced into several isoforms [14], and despite our extensive survey we cannot rule out the possibility of additional unidentified transcripts in *O. fasciatus*.

We designed exon-specific primers to examine the expression of these transcripts using qRT-PCR (table S1). Expression was measured in tissues isolated from teneral adults. Expression of *dsx* transcripts was generally low compared to other genes, and showed considerable variation by sex, tissue and exon (figure 2c). While our primer sets do not allow unambiguous identification of all transcripts, some determinations about transcript-specific expression can be made from these data. Expression of *dsx* was generally higher in females (figure 2c). Transcript isoform D, which contains exon 6, was the most abundant isoform, with particularly strong expression in the ovaries and ovipositor. We also detected this transcript in the male genital capsule, albeit at lower levels. Transcript isoform E, distinguished by exon 2b, was not detected in males, and while it was not strongly expressed in females, isoform E appears to be female-specific. No expression of *dsx* exons was detected in the testes of males, while all exons showed strong expression in the ovaries.

*intersex* and *fruitless* appeared to be expressed strongly and more uniformly across most of the sampled tissues. In *D. melanogaster ix* is required for female somatic development and for sex-specific behavior in females and males [51]. The gene is transcribed in most tissues of both females and males from prepupal to adult stages [52]. High levels of *ix* transcripts were detected in all sampled tissues in both sexes of *O. fasciatus* (figure 2c). Expression was highest in the female ovipositor. *fruitless* is expressed much more strongly in male than in female *D. melanogaster*, particularly in older males [52] where it is required for courtship behavior, fertility, and minor anatomical features specific to males [21–23]. Expression of *fru* in *O. fasciatus* was detected in all sampled tissues, except for the female ovipositor. Expression was strongest in the testes. Interestingly, *fru* expression was comparable and moderate in the heads of females and males (figure 2c).

These patterns of expression for *ix* and *fru* in the milkweed bug are notably less sexually dimorphic than their orthologs in the fruit fly. Excluding the ovipositor, *fru* and *ix* show a strong correlation in expression (Spearman’s rank correlation *rho* = 0.39, *p* = 0.00819). We tested for gene interactions among all three genes by measuring gene expression in an RNA interference (RNAi) background in both sexes. The only such interaction detected was between *ix* and *fru*. Expression of *fru* was significantly reduced in *ix* RNAi males, compared to control males (figure S8; Wilcoxon rank sum test, *p* = 0.019). This implies that *ix* promotes *fru* expression, although this effect may be indirect. The absence of *fru* expression in the ovipositor (figure 3), where *ix* is highly expressed, suggests the action of an unidentified suppressor in that context.

**Figure 3.**
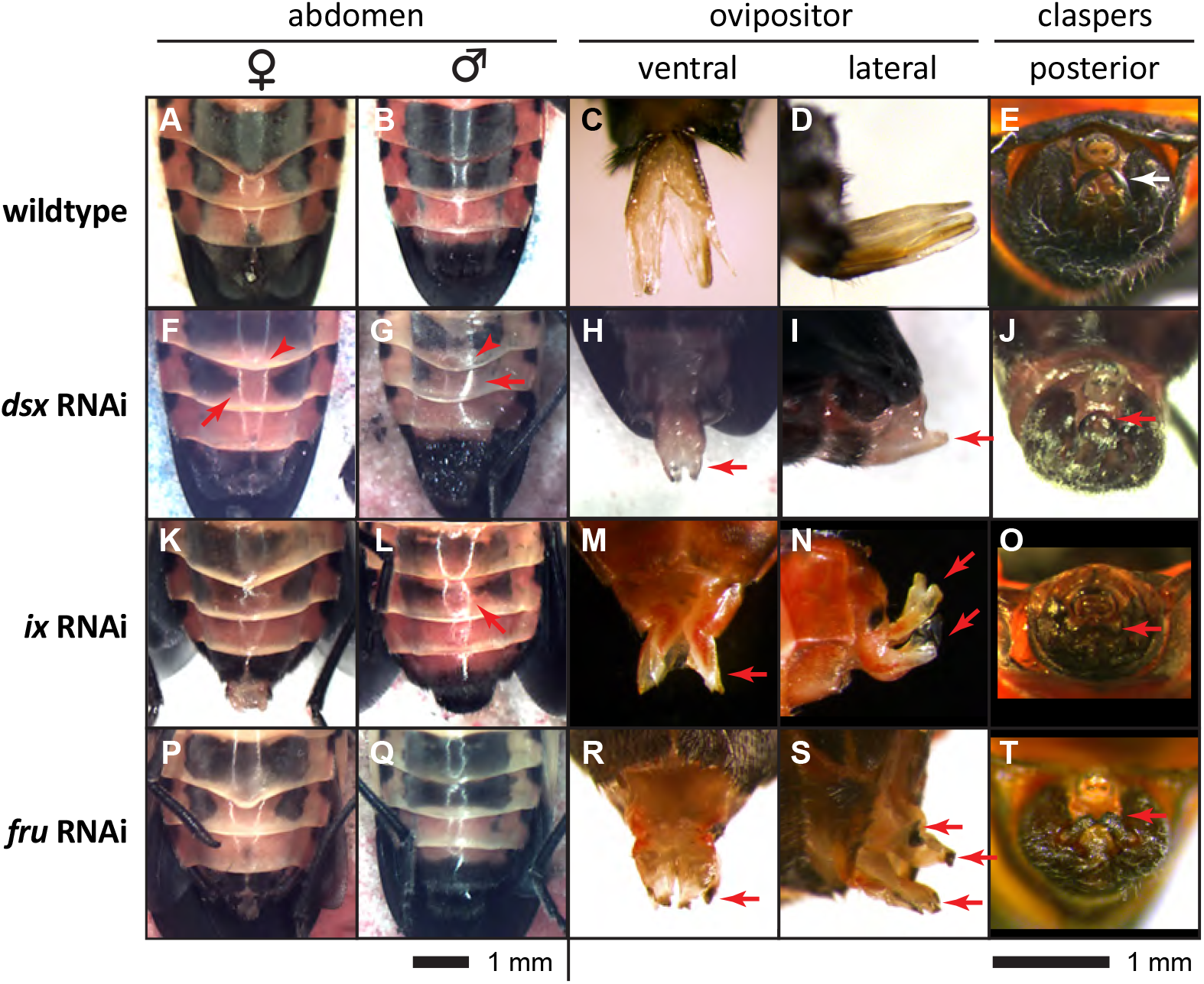
Effects of RNA interference on sexual dimorphisms. (a-e) Unmanipulated *O. fasciatus*. Ventral views of the female and male abdomen differ in the curvature of the fourth sternite, in the distribution of melanic pigment, and in the morphology of the genitalia. Abdomens are shown with anterior towards the top of the page (a,b). The ovipositor is a jointed structure formed by two nested appendages that end in membranous valvulae. Ovipositors are shown from ventral perspective, with anterior up (c), and from left lateral perspective, with dorsal up (d). The male genital capsule (e) is visible from a posterior view; dorsal is up. The posterior of the capsule ends in dorsally pointing claspers (white arrow), used by the male to hold onto females during and after mating. The abdominal panels share the same scale, while the ovipositor and clasper images share a separate scale. One millimeter scale bars are given for each group of panels. Strongly affected RNAi specimens are shown to illustrate phenotypes. Red arrows indicate effected genitalia structures and areas of abdominal pigmentation. (f-j) RNAi targeting *dsx* results in extreme reduction of terminal appendages in both sexes. Only a small nub of tissue represents the valvulae and claspers in severely affected specimens. Abdominal pigmentation develops an intersex pattern in both sexes. Abdominal curvature is also intersex in females and males (red arrowheads). (k-o) *ix* RNAi produces intersex phenotypes in females and males. The female ovipositor is reduced in length and heavily sclerotized, suggesting transformation toward a clasper-like identity. The male genital capsule and claspers are severely reduced. (p-t) *fru* RNAi also reduced the size of the ovipositor and claspers. Claspers are also misshapen, becoming more straight and blunt than in control specimens.

### (c) Development of female and male genitalia requires *ix, fru* and *dsx*

To investigate the developmental functions of *dsx*, *ix* and *fru* in *O. fasciatus*, we performed juvenile-stage RNAi (tables S2, S3). We then examined sexually dimorphic traits in the resulting adults to evaluate loss-of-function phenotypes (figure 3–4). Knockdown of *dsx* significantly affected development of all sexually dimorphic structures in presumptive females and males. Only dsRNAs targeting the exon containing the DNA-binding domain resulted in phenotypic changes (table S3). This exon is present in all transcript isoforms produced from the *dsx* locus (figure 2b). Presumptive female *dsx* RNAi specimens developed ovipositors that were significantly shorter than unmanipulated specimens (figure 4a) and appearing reduced overall, but particularly in the distal valvulae (figure 3h-i). The claspers of presumptive males were also dramatically shorter or absent entirely in some specimens (figure 3j; 5b). Male *dsx* RNAi individuals often died while molting to adulthood, when the genital capsule did not complete molting.

**Figure 4.**
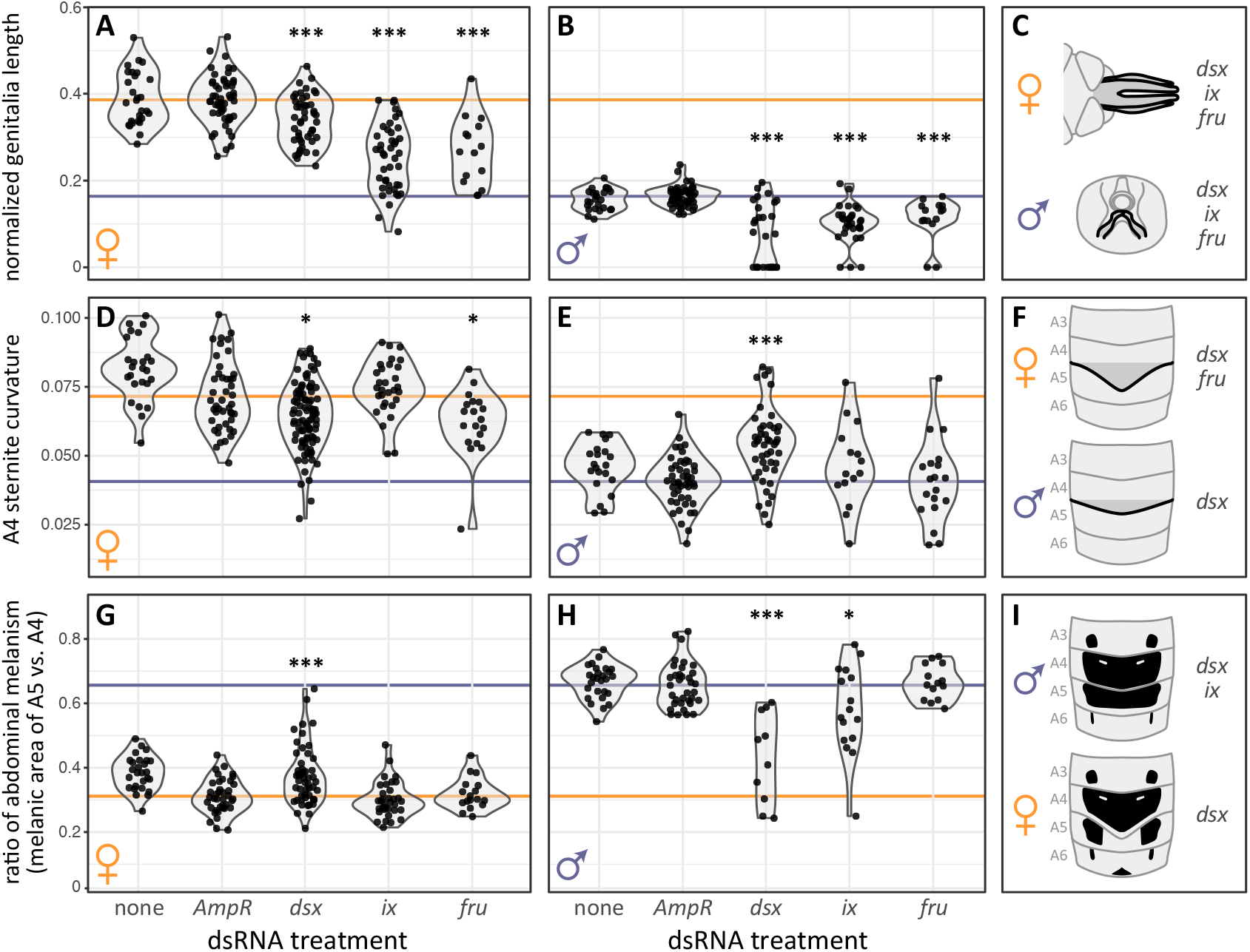
Quantification of RNAi effects on genitalia length (a-b), sternite curvature (d-e) and abdominal melanism (g-h) in females (a,d,g) and males (b,e,h). Dots represent individual specimens. Gray outlines show the relative distribution of values. The mean of control (*AmpR* dsRNA) specimens is indicated by the horizontal line in each panel. Stars indicate significant difference from control values, based on one-sided Wilcoxon rank sum tests with FDR correction. * *p* < 0.05; ** *p* < 0.01; \*\*\**p* < 0.001, and are summarized beside a sketch of typically female and male phenotypes at the right (c,f,i).

In *D. melanogaster* the female isoform of Dsx interacts with the protein encoded by *ix* to activate female-specific target gene expression [20]. Therefore, we tested *ix* function during *O. fasciatus* development. Knockdown of *ix* by RNAi produced intersex phenotypes (figure 3k-o; 5a-b,h), although it was still possible to identify bugs unambiguously as female and male based on the number of terminal appendages and the presence of a genital capsule in presumptive males. Length was significantly reduced in ovipositors and claspers of *ix* RNAi specimens (figure 4a-b). Distally the ovipositor developed with heavy sclerotization, suggesting partial transformation toward a clasper-like identity (figure 3m-n). In presumptive males, the genital capsule and claspers were significantly reduced in size (figure 3o; 4b).

The transcription factor encoded by *fru* is conserved among insects (figure S4). In *D. melanogaster fru* is required for male-specific behaviors, but plays a relatively minor role in anatomical development [48]. Surprisingly, knockdown of *fru* in *O. fasciatus* caused dramatic defects in sexually dimorphic adult structures (figure 3p-t), similar in many ways to the effects of *dsx* and *ix* knockdowns. *fru* RNAi significantly reduced the length of the ovipositor and claspers (figure 4a-b). While the ovipositors of *ix* RNAi specimens decreased in length, they also increased in thickness and pigmentation, suggestive of partial transformation to clasper identity. In contrast, the ovipositors of *fru* RNAi specimens did not appear transformed, and retained a membranous structure (compare figure 3m-n to 3r-s). Some presumptive female *fru* RNAi specimens also showed necrosis of the distal valvulae (figure 3s). In *fru* RNAi males, claspers were reduced in size (figure 4b) and often misshapen, lacking their characteristic arch (figure 3t). Expression of *fru* in the ovipositor of teneral adults was very low (figure 2c), suggesting that *fru* is required at an earlier stage of ovipositor development or that it acts indirectly by activating factors that induce ovipositor or genitalia development.

Other appendages in *dsx*, *ix* and *fru* RNAi individuals, such as the antennae, legs and wings were not obviously reduced in length or otherwise different from control specimens. While RNAi targeting *dsx*, *ix* and *fru* produced dramatic phenotypes in the genitalia, we did not identify obvious knockdown phenotypes in the testes or ovaries of RNAi specimens (figure S7). However, we could not rule out the possibility of smaller scale anatomical changes.

### (d) *dsx* inhibits intersex sternite shapes in females and in males

One of the most obvious sexual dimorphisms in milkweed bugs is the posterior projection of the fourth abdominal sternite (figure 1a). We examined the effects of RNAi gene knockdown on sternite curvature, quantified using nine landmarks placed on the sternite’s posterior edge (figure 1a-b). After Procrustes alignment of the coordinate positions, these landmarks were regressed to a line. The square-root of the mean of squared residuals (residual mean standard deviation) was used as a metric of curvature. Sternite curvature is not a binary trait in milkweed bugs. Females and males display a continuous range of curvature (figure 4d-f). Nevertheless, among dsRNA control specimens, the sternites of females and males showed significantly different median curvatures (Wilcoxon rank sum test, *p* < 2.2×10^-16^).

Male milkweed bugs retain a juvenile sternite shape, while the female sternite projection arises during the last two molts before adulthood. Therefore, we predicted that inhibiting genes required for the development of sexual dimorphism might cause presumptive females to also retain the juvenile sternite shape. Surprisingly, *dsx* RNAi produced sternite shape changes in female and in male bugs. As expected for females, sternite curvature in *dsx* RNAi presumptive females was significantly reduced compared to that of dsRNA control females (figure 4d; Wilcoxon rank sum test, *p* = 4.73×10^-3^). However, among presumptive male *dsx* RNAi specimens, sternite curvature increased (*p* = 1.12×10^-7^), adopting a more typically female phenotype. A similar effect of less magnitude was also produced by the simultaneous knockdown of all three *doublesex* paralogs (females: *p* = 0.0458; males: *p* = 4.15×10^-3^). Thus, for both sexes, *dsx* RNAi caused sternite shapes to become more intermediate.

### (e) Development of female sternite shape requires *fru*

Since knock-down of *ix* and *fru* produced dramatic phenotypes in the genitalia of female and male milkweed bugs, we also examined other sexually dimorphic traits in these specimens. RNAi targeting *ix* did not significantly affect the sternite curvature (figure 4d-e). Even specimens with dramatic intersex genitalia still displayed sternite curvatures typical of their presumptive sex (figure S9). In contrast, knockdown of *fru* reduced the curvature of sternites in females (*p* = 0.0107). While some presumptive female *fru* RNAi specimens had sternites with a typically male shape, most developed with an intermediate curvature (figure 4d). However, presumptive male *fru* RNAi specimens were not different from dsRNA control specimens (figure 4e; *p* = 0.831). This suggests that *fru* activity is required for the increased curvature of the A4 sternite in females, but not in males, which retain a more juvenile sternite shape.

### (f) Sex-specific distribution of abdominal melanism requires *dsx*

The ventral abdomen of *O. fasciatus* typically has several areas of melanic pigmentation. While considerable individual variation exists, females and males have distinct (non-overlapping) ratios of melanism on the fifth and fourth abdominal segments (figure 4g-i). Females have less melanism on A5, compared to males where melanic pigmentation is more similar on A4 and A5. Within each sex, this trait is independent of the curvature of the A4 sternite (figure S10; Spearman’s rank correlation for females *p* = 0.72; for males *p* = 0.22).

RNA interference targeting *dsx* affected the distribution of abdominal melanism in both sexes (figure 3f-g, 5g-i). Females normally have much less melanic pigmentation on the fifth abdominal sternite, compared to the fourth. However, presumptive female *dsx* RNAi specimens had significantly more even distribution of melanism than control females (Wilcoxon rank sum test, *p* = 2.19×10^-4^). Conversely, presumptive male *dsx* RNAi specimens had a reduction in the ratio of A5 to A4 melanic area, compared to male *AmpR* RNAi specimens (*p* = 7.62×10^-7^). In contrast to control specimens, where the ratio of abdominal pigmentation is completely non-overlapping, there is no longer any sexual dimorphism in this trait among *dsx* RNAi specimens (*p* = 0.25).

### (g) *ix* is required in males for sex-specific abdominal melanism

As with sternite shape, we also examined abdominal pigmentation following RNAi targeting *ix* and *fru*. While *fru* RNAi produced significant changes in the sternite shapes of presumptive females (figure 4d), these specimens did not differ significantly from controls in their distribution of abdominal melanism (figure 4g; *p* = 0.41). RNAi targeting *ix* did affect abdominal pigmentation, but only in presumptive males. Often presumptive male *ix* RNAi specimens had reduced pigmentation on the fifth abdominal segment, leading to a more typically female pigmentation pattern (figure 4h) that was significantly less evenly distributed than in control male bugs (*p* = 0.0173).

## 4. Discussion

### (a) Sexual dimorphism requires *dsx*

Our results suggest a general role for *dsx* in the development of adult sexual dimorphisms in *O. fasciatus*. All three of the sexually dimorphic traits examined in this study require *dsx* function. Interestingly, the function of *dsx* was not simply to promote the development of sexually dimorphic traits. Genitalia are the most obviously dimorphic structures differing in females and males, and in these appendages *dsx* RNAi reduced their length in all individuals (figure 4a-b). While these effects could be interpreted as a generalized role in promoting growth, *dsx* RNAi did not effect other appendages and a more switch-like role appears for *dsx* in the development of other sexually dimorphic structures. RNAi knockdown of *dsx* produced more female-like sternite curvature in males and more male-like curvature in females, suggesting that *dsx* acts in each sex to inhibit intersex phenotypes, just as it does in *Drosophila* [53]. In effect, *dsx* makes the distribution of sternite curvature more bimodal. Each of these traits develops from a null state in juveniles. Males retain the relatively uncurved juvenile sternite shape, while neither sex has abdominal melanism as a juvenile. If the role of *dsx* was to promote sternite curvature via growth promotion alone, then we would expect male *dsx* RNAi specimens to be comparable to control males. Similarly, if *dsx* promoted abdominal pigmentation generally, we might expect reduced pigmentation overall with no significant effect on its distribution. However, both these traits are more intersex in their values following *dsx* RNAi. It is likely that distinct isoforms of Dsx act on distinct target genes to produce different effects in each sex, and that these intersex phenotypes arise from the absence of this regulation.

Expression of *dsx* in *O. fasciatus* was strongly female-biased and one *dsx* transcript (isoform E) appeared only in females (figure 2c). Similarly, in the sweet potato whitefly, *Bemisia tabaci*, a single *dsx* ortholog also has sex-biased expression [54]. Although in that member of the Sternorrhyncha, *dsx* is more strongly expressed in males and RNAi knockdown revealed a requirement in male genitalia development, and merely a role in vitellogenin regulation in females. In contrast, *dsx* in *O. fasciatus* was expressed strongly in ovaries, but not at all in testes. While gonads of RNAi specimens did not display anatomical defects (figure S7), these expression patterns suggest that sex-specific divergence of transcripts may play a role in the evolution of sex-specific development.

### (b) Sexual dimorphisms develop by distinct mechanisms in milkweed bugs

Both female and male *O. fasciatus* require *ix* and *fru* for proper development, but in different sexually dimorphic traits (figure 4c,f,i). Orthologs of these genes in *D*. *melanogaster* have much more limited roles, affecting only one sex or the other. Interestingly, while *ix* and *fru* affect genitalia development in both sexes of *O. fasciatus*, these genes affect different sexually dimorphic traits, in different sexes.

*fruitless* RNAi only affected the development of female, but not male, sternite curvature (figure 4d-f). Because male sternite curvature closely resembles the phenotype of juveniles, it is possible that *fru* is involved in regulating growth and proliferation of these cells, rather than in directly determining sexually dimorphic identity, as seems to be the case for *dsx*. Knockdown of *ix* did not significantly change sternite curvature, but affected the distribution of abdominal pigmentation only in male milkweed bugs (figure 4g-i). It is possible that only some Dsx isoforms in *O. fasciatus* interact with Ix, in a trait-specific context, rather than in a sex-specific context. Isoform D of *O. fasciatus dsx* is a strong candidate for this interaction, because of its high expression in both female and male genitalia, but lower expression in other somatic body regions (figure 3). Confirmation of this hypothesis must await the development of isoform-specific antibodies from milkweed bug Dsx. Additionally, it is possible that the expression of distinct *dsx* isoforms is regulated by tissue-specific enhancers. A recent study of regulatory sequences in *D. melanogaster* found that expression of *dsx* in distinct sexually dimorphic structures is controlled by separate enhancers [55]. Similarly, analysis of gene expression in different larval and pupal tissues of the butterfly *Papilio polytes* has suggested distinct tissue-specific targets of *dsx* activity [56].

These findings support the concept that some aspects of sex determination in insects may be structure-specific. It is possible that this form of regulation may be an evolutionary intermediate. If a gene has multiple enhancers for expression in different areas, as exemplified by *D. melanogaster dsx*, then a mutation eliminating one enhancer would produce the kind of mosaic genetic requirement seen for *O. fasciatus ix* and *fru*.

### (c) Evolution of sex determination mechanisms in insects

The results we present here further illuminate the diversity of sex determination mechanisms in insects. In the crustacean *Daphnia magna* [33] and in the cockroach *Blattella germanica* [32] *dsx* is required for male development but not for female development. In Holometabola, such as *Drosophila*, females and males require sex-specific isoforms of *dsx* for sexually dimorphic development [53]. The evolutionary timing of this transition remains uncertain. RNA interference of *dsx* in another hemipteran, the brown planthopper *Nilaparvata lugens*, produces a pseudo-female phenotype in presumptive males, but comparably mild phenotypes in females, which remain fertile [31]. This could be viewed as an intermediate state, suggesting that the transition from male-specific *dsx* function to a functional requirement for sex-specific isoforms, began at the time that Polyneoptera, such as *B. germanica*, diverged from other insects, about 390 Mya [37], and continued in the lineage leading to Holometabola, which split from Hemiptera around 375 Mya [37]. The finding that *O. fasciatus* requires *dsx* activity in females and males suggests instead that the evolution of *dsx* function may be much more dynamic, requiring either independent gains of *dsx* activity in female development in *O. fasciatus* and Holometabola, or that *dsx* was involved in female development after Hemiptera and Holometabola split from Polyneoptera and that female *dsx* function was then lost in the lineage leading to *B. germanica*.

Compared to other genes in insects involved in sexually dimorphic development, such as *ix* and *fru, doublesex* orthologs evolve rapidly in their protein sequence [26]. This could suggest that *dsx* occupies a space in the developmental network where mutations are more readily tolerated [24]. Other genes, including *ix* and *fru*, may be more limited in the evolution of their protein sequences by stabilizing selection, perhaps due to greater pleiotropy. However, differences in the functions of these genes among insect species suggest that regulatory changes still occur without comparable constraints.

Sex determination pathways differ substantially among animals [1–3], evolving by changes gene expression patterns, splicing, and the interactions of pathway proteins [3,25,27]. The functions of *ix* and *fru* in *O. fasciatus* are different from those reported in other insects. In *D. melanogaster, ix* is only required in females for genitalia development while both female and male milkweed bugs require *ix* for genitalia development (figure 3–4) [36]. *ix* RNAi knockdown in *N. lugens* produced genitalia defects in females, but not in males [57]. *fru* is a transcription factor required for development of male-specific neurons in *D. melanogaster* [14]. In *O. fasciatus*, we find a more expansive requirement for *fru* in female and male genitalia development (figure 3r-t; 4a-c) and in the development of female sternite curvature (figure 4d,f). To the best of our knowledge, such a prominent function for *fru* has not been previously reported. Further studies of *fru* outside drosophilids are needed to better understand its functional diversity across the insects and to determine when during drosophilid evolution it became more restricted in its developmental role. Overall, these differences in the functions of *ix* and *fru* suggest that somatic sex determination may have a much greater diversity of mechanisms among insects than is currently understood.

### (d) Conclusions

In insects such as the fruit fly, *dsx* functions as a critical regulator of somatic sex determination for females and males [2,4], while outside Holometabola *dsx* has been found necessary for male development primarily [31] or entirely [32, 33]. We find that both sexes in *O. fasciatus* require *dsx* activity in multiple dimorphic traits (figures 3–4). Knockdowns of *dsx* or all three DMRT genes together (figure S11) during juvenile development do not completely eliminate adult sexual dimorphism. Rather, *dsx*, together with *ix* and *fru*, appears to help facilitate trait-specific development of sexual dimorphism, including the development of female and male genitalia and the inhibition of intersex values for sternite curvature and abdominal pigmentation of both sexes. Nevertheless, it is likely that unidentified genes play a proximate role in milkweed bug sex determination. More generally, these results show that genes may act to control the development of somatic sexual dimorphisms by acting in tissue-specific as well as sex-specific contexts. The sex determination cascades of other insects should be investigated in more detail to expand our understanding of the evolution and diversity of sex determination mechanisms. Such studies will broaden our perspective on developmental pathway evolution and help us better understand how genetic networks bias evolutionary outcomes.

## Data accessibility

DNA sequences were archived in GenBank (accession numbers MZ197788 - MZ197800). All other data were archived at Dryad (https://doi.org/10.5061/dryad.kwh70rz3q), including amino acid sequence alignments of candidate gene orthologs and resulting Newick tree files, tables of morphological measurements and 2D landmark coordinate positions for sternite shapes, gene expression data, and the R scripts used for analysis.

## Author contributions

JJ and ML designed and conducted most experiments and contributed equally to data analysis, creation of draft figures, and writing of the main text. YJL, MY, and ZZ conducted portions of the experiments and contributed to data analysis. DRA designed and supervised all experiments, secured funding, prepared final figures, and contributed to data analysis and to the writing of the main text. All authors gave final approval for publication and agreed to be held accountable for the work performed therein.

## Competing interests

We declare we have no competing interests.

## Funding

Research reported in this publication was supported by the Colby College Division of Natural Sciences, by an Institutional Development Award (IDeA) from the National Institute of General Medical Sciences of the National Institutes of Health under grant number P20GM0103423, and by grant IOS-1350207 from the National Science Foundation to DRA.

## Acknowledgements

The authors wish to thank several people for their support and comments on an early draft of the manuscript: Devin M. O’Brien, Joshua L. Steele, Lyndell M. Bade, Johanna van Oers, Russell Johnson, David L. Stern, and Cassandra G. Extavour. Special thanks to Kristen Panfilio for sharing transcriptome data prior to their publication.

## Notes

https://doi.org/10.5061/dryad.kwh70rz3q

### Competing Interest Statement

The authors have declared no competing interest.

### Summary of Updates

The three genes described as doublesex paralogs in the initial manuscript were found to be orthologs of three of the DMRT family genes, known from other animals. The discussion of these genes has been changed to reflect that orthology.

## References

1. Schütt C, Nöthiger R. 2000 Structure, function and evolution of sex-determining systems in Dipteran insects. Development 127, 667–677.

2. Kopp A. 2012 Dmrt genes in the development and evolution of sexual dimorphism. Trends Genet. 28, 175–184. (doi:10.1016/j.tig.2012.02.002)

3. Herpin A, Schartl M. 2015 Plasticity of gene-regulatory networks controlling sex determination: of masters, slaves, usual suspects, newcomers, and usurpators. EMBO Rep. 16, 1260–1274. (doi:10.15252/embr.201540667)

4. Arbeitman MN, Fleming AA, Siegal ML, Null BH, Baker BS. 2004 A genomic analysis of *Drosophila* somatic sexual differentiation and its regulation. Development 131, 2007–2021. (doi:10.1242/dev.01077)

5. Rideout EJ, Dornan AJ, Neville MC, Eadie S, Goodwin SF. 2010 Control of sexual differentiation and behavior by the doublesex gene in *Drosophila melanogaster*. Nat. Neurosci. 13, 458–466. (doi:10.1038/nn.2515)

6. Robinett CC, Vaughan AG, Knapp J-M, Baker BS. 2010 Sex and the Single Cell. II. There Is a Time and Place for Sex. PLoS Biol. 8, e1000365. (doi:10.1371/journal.pbio.1000365)

7. Tanaka K, Barmina O, Sanders LE, Arbeitman MN, Kopp A. 2011 Evolution of Sex-Specific Traits through Changes in HOX-Dependent doublesex Expression. PLoS Biol. 9, e1001131. (doi:10.1371/journal.pbio.1001131)

8. Kiuchi T et al. 2014 A single female-specific piRNA is the primary determiner of sex in the silkworm. Nature 509, 633–636. (doi:10.1038/nature13315)

9. Sharma A et al. 2017 Male sex in houseflies is determined by *Mdmd*, a paralog of the generic splice factor gene *CWC22*. Science 356, 642–645. (doi:10.1126/SCIENCE.AAM5498)

10. Baker BS, Ridge KA. 1980 Sex and the single cell. I. On the action of major loci affecting sex determination in *Drosophila melanogaster*. Genetics 94, 383–423.

11. Erickson JW, Quintero JJ. 2007 Indirect Effects of Ploidy Suggest X Chromosome Dose, Not the X:A Ratio, Signals Sex in *Drosophila*. PLOS Biol. 5, e332. (doi:10.1371/JOURNAL.PBIO.0050332)

12. Butler B, Pirrotta V, Irminger-Finger I, Nöthiger R. 1986 The sex-determining gene *tra* of *Drosophila*: molecular cloning and transformation studies. EMBO J. 5, 3607–3613. (doi:10.1002/j.1460-2075.1986.tb04689.x)

13. McKeown M. 1992 Sex differentiation: The role of alternative splicing. Curr. Opin. Genet. Dev. 2, 299–303. (doi:10.1016/S0959-437X(05)80288-6)

14. Demir E, Dickson BJ. 2005 *fruitless* splicing specifies male courtship behavior in *Drosophila*. Cell 121, 785–794. (doi:10.1016/j.cell.2005.04.027)

15. Boggs RT, Gregor P, Idriss S, Belote JM, McKeown M. 1987 Regulation of sexual differentiation in *D. melanogaster* via alternative splicing of RNA from the *transformer* gene. Cell 50, 739–747. (doi:10.1016/0092-8674(87)90332-1)

16. Inoue K, Hoshijima K, Sakamoto H, Shimura Y. 1990 Binding of the *Drosophila Sex-lethal* gene product to the alternative splice site of *transformer* primary transcript. Nature 344, 461–463. (doi:10.1038/344461a0)

17. Burtis KC, Baker BS. 1989 *Drosophila doublesex* gene controls somatic sexual differentiation by producing alternatively spliced mRNAs encoding related sex-specific polypeptides. Cell 56, 997–1010. (doi:10.1016/0092-8674(89)90633-8)

18. Tian M, Maniatis T. 1993 A splicing enhancer complex controls alternative splicing of *doublesex* pre-mRNA. Cell 74, 105–114. (doi:10.1016/0092-8674(93)90298-5)

19. Waterbury JA, Jackson LL, Schedl P. 1999 Analysis of the doublesex female protein in *Drosophila melanogaster*: role on sexual differentiation and behavior and dependence on *intersex*. Genetics 152, 1653. (doi:10.1093/genetics/152.4.1653)

20. Garrett-Engele CM, Siegal ML, Manoli DS, Williams BC, Li H, Baker BS. 2002 *intersex*, a gene required for female sexual development in *Drosophila*, is expressed in both sexes and functions together with *doublesex* to regulate terminal differentiation. Development 129, 4661–4675. (https://doi.org/10.1242/dev.129.20.4661)

21. Hall J. 1978 Courtship among males due to a male-sterile mutation in *Drosophila melanogaster*. Behav. Genet. 8, 125–141. (doi:10.1007/BF01066870)

22. Hall JC. 1994 The mating of a fly. Science 264, 1702–1714. (doi:10.1126/science.8209251)

23. Usui-Aoki K et al. 2000 Formation of the male-specific muscle in female *Drosophila* by ectopic fruitless expression. Nat. Cell Biol. 2, 500–506. (doi:10.1038/35019537)

24. Stern DL, Orgogozo V. 2009 Is genetic evolution predictable? Science 323, 746–51. (doi:10.1126/science.1158997)

25. Salz HK. 2011 Sex determination in insects: A binary decision based on alternative splicing. Curr. Opin. Genet. Dev. 21, 395–400. (doi:10.1016/j.gde.2011.03.001)

26. Laslo M, Just J, Angelini DR. 2022 Theme and variation in the evolution of insect sex determination. Journal of Experimental Zoology. Part B, Molecular and Developmental Evolution.In revision.

27. Shukla JN, Nagaraju J. 2010 Doublesex: A conserved downstream gene controlled by diverse upstream regulators. J. Genet. 89, 341–356. (doi:10.1007/s12041-010-0046-6)

28. Shukla JN, Palli SR. 2012 Doublesex target genes in the red flour beetle, *Tribolium castaneum*. Sci. Reports 2012 21 2, 1–10. (doi:10.1038/srep00948)

29. Verhulst EC, van de Zande L. 2015 Double nexus—Doublesex is the connecting element in sex determination. Brief. Funct. Genomics 14, 396–406. (doi:10.1093/BFGP/ELV005)

30. Geuverink E, Rensink AH, Rondeel I, Beukeboom LW, van de Zande L, Verhulst EC. 2017 Maternal provision of *transformer-2* is required for female development and embryo viability in the wasp *Nasonia vitripennis*. Insect Biochem. Mol. Biol. 90, 23–33. (doi:10.1016/J.IBMB.2017.09.007)

31. Zhuo JC, Hu QL, Zhang HH, Zhang MQ, Jo SB, Zhang CX. 2018 Identification and functional analysis of the *doublesex* gene in the sexual development of a hemimetabolous insect, the brown planthopper. Insect Biochem. Mol. Biol. 102, 31–42. (doi:10.1016/j.ibmb.2018.09.007)

32. Wexler J, Delaney EK, Belles X, Schal C, Wada-Katsumata A, Amicucci MJ, Kopp A. 2019 Hemimetabolous insects elucidate the origin of sexual development via alternative splicing. Elife 8. (doi:10.7554/ELIFE.47490)

33. Kato Y, Kobayashi K, Watanabe H, Iguchi T. 2011 Environmental sex determination in the branchiopod crustacean *Daphnia magna*: Deep conservation of a *doublesex* gene in the sex-determining pathway. PLOS Genet. 7, e1001345. (doi:10.1371/JOURNAL.PGEN.1001345)

34. Clynen E, Ciudad L, Bellés X, Piulachs M-D. 2011 Conservation of *fruitless*’ role as master regulator of male courtship behaviour from cockroaches to flies. Dev. Genes Evol. 2011 2211 221, 43–48. (doi:10.1007/S00427-011-0352-X)

35. Yang X, Chen K, Wang Y, Yang D, Huang Y. 2021 The sex determination cascade in the silkworm. Genes 12, 315. (doi:10.3390/genes12020315)

36. Aspiras AC, Smith FW, Angelini DR. 2011 Sex-specific gene interactions in the patterning of insect genitalia. Dev. Biol. 360, 369–380. (doi:10.1016/j.ydbio.2011.09.026)

37. Misof B, Liu S, Meusemann K, Peters RS, Al. E. 2014 Phylogenomics resolves the timing and pattern of insect evolution. Science 346, 763–767. (doi:10.1017/CBO9781107415324.004)

38. Liu P, Kaufman TC. 2009 Morphology and Husbandry of the Large Milkweed Bug, *Oncopeltus fasciatus*. Cold Spring Harb. Protoc. 2009, pdb.emo127--. (doi:10.1101/pdb.emo127)

39. Ewen-Campen B, Shaner N, Panfilio K, Suzuki Y, Roth S, Extavour CG. 2011 The maternal and early embryonic transcriptome of the milkweed bug *Oncopeltus fasciatus*. BMC Genomics 12, 61. (doi:10.1186/1471-2164-12-61)

40. Panfilio KA et al. 2019 Molecular evolutionary trends and feeding ecology diversification in the Hemiptera, anchored by the milkweed bug genome. Genome Biol. 20, 64. (doi:10.1186/s13059-019-1660-0)

41. Sievers F et al. 2011 Fast, scalable generation of high-quality protein multiple sequence alignments using Clustal Omega. Mol. Syst. Biol. 7, 539. (doi:10.1038/MSB.2011.75)

42. Stamatakis A. 2014 RAxML version 8: a tool for phylogenetic analysis and post-analysis of large phylogenies. Bioinformatics 30, 1312–1313. (do:10.1093/bioinformatics/btu033)

43. Su J, Zhang R, Dong J, Yang C. 2011 Evaluation of internal control genes for qRT-PCR normalization in tissues and cell culture for antiviral studies of grass carp (*Ctenopharyngodon idella*). Fish Shellfish Immunol. 30, 830–835. (doi:10.1016/J.FSI.2011.01.006)

44. Schneider CA, Rasband WS, Eliceiri KW. 2012 NIH Image to ImageJ: 25 years of image analysis. Nat. Methods 9, 671–675. (doi:10.1038/nmeth.2089).

45. Rohlf FJ. 2015 The tps series of software. Hystrix 26, 9–12. (https://doi.org/10.4404/hystrix-26.1-11264)

46. Adams DC, Collyer ML, Kaliontzopoulou A. 2020 Geomorph: Software for geometric morphometric analyses. R package version 3.2.1. (https://cran.r-project.org/package=geomorph)

47. Sharma AI, Yanes KO, Jin L, Garvey SL, Taha SM, Suzuki Y. 2016 The phenotypic plasticity of developmental modules. Evodevo 7, 1–14. (doi:10.1186/S13227-016-0053-7/FIGURES/8)

48. Ryner LC, Goodwin SF, Castrillon DH, Anand A, Villella A, Baker BS, Hall JC, Taylor BJ, Wasserman SA. 1996 Control of male sexual behavior and sexual orientation in *Drosophila* by the *fruitless* gene. Cell 87, 1079–1089. (doi:10.1016/S0092-8674(00)81802-4)

49. Brunner B et al. 2001 Genomic organization and expression of the *doublesex*-related gene cluster in vertebrates and detection of putative regulatory regions for *DMRT1*. Genomics 77, 8–17. (doi:10.1006/GENO.2001.6615)

50. Price DC, Egizi A, Fonseca DM. 2015 The ubiquity and ancestry of insect *doublesex*. Sci. Rep. 5, 1–9. (doi:10.1038/srep13068)

51. Nagoshi RN, McKeown M, Burtis KC, Belote JM, Baker BS. 1988 The control of alternative splicing at genes regulating sexual differentiation in *D. melanogaster*. Cell 53, 229–236. (doi:10.1016/0092-8674(88)90384-4)

52. Matthews BB et al. 2015 Gene model annotations for *Drosophila melanogaster*: Impact of high-throughput data. G3 Genes, Genomes, Genet. 5, 1721–1736. (doi:10.1534/G3.115.018929/-/DC1)

53. Hildreth PE. 1965 *Doublesex*, a recessive gene that transforms both males and females of *Drosophila* into intersexes. Genetics 51, 659. (doi:10.1093/genetics/51.4.659)

54. Guo L, Xie W, Liu Y, Yang Z, Yang X, Xia J, Wang S, Wu Q, Zhang Y. 2018 Identification and characterization of doublesex in *Bemisia tabaci*. Insect Mol. Biol. 27, 620–632. (doi:10.1111/imb.12494)

55. Rice GR, Barmina O, Luecke D, Hu K, Arbeitman M, Kopp A. 2019 Modular tissue-specific regulation of *doublesex* underpins sexually dimorphic development in *Drosophila*. Development 146. (doi:10.1242/dev.178285)

56. Deshmukh R, Lakhe D, Kunte K. 2020 Tissue-specific developmental regulation and isoform usage underlie the role of *doublesex* in sex differentiation and mimicry in *Papilio* swallowtails. R. Soc. Open Sci. 7, 200792. (doi:10.1098/rsos.200792)

57. Zhang H-H, Xie Y-C, Li H-J, Zhuo J-C, Zhang C-X. 2021 Pleiotropic roles of the orthologue of the *Drosophila melanogaster intersex* gene in the brown planthopper. Genes 12, 379. (doi:10.3390/genes12030379)

