## Supplemental Tables and Figures for "Distinct developmental mechanisms influence sexual dimorphisms in the milkweed bug *Oncopeltus fasciatus*"

### Supplemental Information

**Supplemental Table 1.** Primer and probe sequences used for real-time PCR.

| target | *exon* | primer name | sequence | position | amplicon (bp) | reporter | quencher |
|---|---|---|---|---|---|---|---|
| *EF1a* | | Of-EF1a-396P | AACCAGAGAGCATGCCCTTC | 396 | 130 | TxRed | BHQ2 |
| | | Of-EF1a-358F | ACTGGTGAGTTTGAAGCTGGTAT | 358 | | | |
| | | Of-EF1a-487R | AGTAGGGTGGTTCAGTTGAATCC | 464 | | | |
| *ix* | | Of-ix-476P | TAGCCCCGACGAGAAGTGAC | 476 | 126 | Cy5.5 | BHQ3 |
| | | Of-ix-452F | GTCAGCGCTATCTATCTTTGGGA | 452 | | | |
| | | Of-ix-577R | CCTTAGCAAAAGTGACCTGTGTG | 555 | | | |
| *fru* | | Of-fru-424P | CAAATTCGTGGCTTGGCTGA | 424 | 90 | Cy5.5 | BHQ3 |
| | | Of-fru-379F | AAGCAAGATGAACTAGCTGGGAT | 379 | | | |
| | | Of-fru-468R | ATCTTGTCTAGGTGCTTCACGTG | 446 | | | |
| *dsx* | 1 | Of-dsxc-183P | CGTAAAGAAACGTCGTGCCG | 183 | 126 | HEX | BHQ1 |
| | | Of-dsxc-100F | CACAAGAATGACTGTCCGTTCAC | 100 | | | |
| | | Of-dsxc-225R | AGCAAGTTCTTCCTGGAGTGAAT | 202 | | | |
| | 2 | Of-dsxc-314P | TGAAAGAAGTAGTCAGCAAGAAGC | 314 | 77 | FAM | BHQ1 |
| | | Of-dsxc-290F | CGAAACAACCCTCCATCAACAAG | 290 | | | |
| | | Of-dsxc-366R | CGTCGGAGCTTTCTGCAGTATAT | 344 | | | |
| | 2b | Of-dsxc-427P | TTGCTTCTAGCCCTGGGATC | 427 | 71 | FAM | BHQ1 |
| | | Of-dsxc-399F | TAAGCACTGTTCGAAACACTCAC | 399 | | | |
| | | Of-dsxc-469R | TATCAGACAACTACGGAAAGGGC | 447 | | | |
| | 3 | Of-dsxc-1203P | AAAGAGCGGTTGTCCAGAAA | 1203 | 82 | FAM | BHQ1 |
| | | Of-dsxc-1175F | GGACAGAAATAGACAACTGGCTT | 1175 | | | |
| | | Of-dsxc-1256R | AGTAAAGTATCACAGGCTCTCGG | 1234 | | | |
| | 4 | Of-dsxc-1500P | CAACCGGCCCTCGACAGG | 1500 | 64 | FAM | BHQ1 |
| | | Of-dsxc-1477F | CCAGTGAGCTGTGTGAAGAACTT | 1477 | | | |
| | | Of-dsxc-1540R | GAGATAAGACGGAGGCCATCTTC | 1518 | | | |
| | 6 | Of-dsxc-1745P | ACAAGTATTTCGTGTTGTACAGCA | 1745 | 164 | FAM | BHQ1 |
| | | Of-dsxc-1647F | CCCCAATTAATCTGGTATTGAAAATTAA | 1647 | | | |
| | | Of-dsxc-1810R | ACGAAATTTGCCGTAATATTGAAAAG | 1785 | | | |
| *dmrt99B* | 1 | Of-dsxa-158P | AGCAGGCACAGGAGGAGAG | 158 | 112 | Cy5 | BHQ3 |
| | | Of-dsxa-95F | CCAAGTGTACCCTCATCGCC | 95 | | | |
| | | Of-dsxa-206R | GAGTACACTAGCCCGAGTTCC | 186 | | | |
| | 2 | Of-dsxa-362P | CGGGACACGGAGAGGAGG | 362 | 64 | FAM | BHQ1 |
| | | Of-dsxa-339F | CCTTCAAATTGACAGAGGACAGC | 339 | | | |
| | | Of-dsxa-402R | GGAGACATCTTCAGGTTCCACTT | 380 | | | |
| | 3 | Of-Dsx-768P | CGAAAACACAGAAGAAACACCCC | 768 | 74 | FAM | BHQ1 |
| | | Of-Dsx-743F | GTAAGAGGGCAAGACTCTCGGAG | 743 | | | |
| | | Of-Dsx-816R | AATGATGGGTTGTGCTGGAATTG | 794 | | | |
| | 5 | Of-dsxa-1238P | TCTAGCCCGGCGTCCTAC | 1238 | 83 | FAM | BHQ1 |
| | | Of-dsxa-1208F | GGACCTCGTATTCCGAATGACTG | 1208 | | | |
| | | Of-dsxa-1290R | TACGGATCAGAATAAGTCAGCCG | 1268 | | | |

| | | | | | | | |
|---|---|---|---|---|---|---|---|
| *dmrt93B* | 1 | Of-dsxb-79P | AAGGGCACAAGAAGGACTGC | 79 | 99 | FAM | BHQ1 |
| | | Of-dsxb-53F | GAACCATGGGATCATCAGCTGG | 53 | | | |
| | | Of-Dxx-133R | CTTTGCCTTTCAGCGATAAGGAC | 111 | | | |
| | 3 | Of-Dxx-399P | CTCGTCTTCAACAGGTGTGACG | 399 | 102 | FAM | BHQ1 |
| | | Of-Dxx-362F | GAAATTGTTCCCAGACAGGAAGC | 362 | | | |
| | | Of-Dxx-463R | TTGAAGGAAGACAGACAGAGCTG | 441 | | | |
| | 4a | Of-Dxx-467P | GGTTATGAACTGAAACCGGCCT | 467 | 71 | FAM | BHQ1 |
| | | Of-Dxx-441F | CAGCTCTGTCTGTCTTCCTTCAA | 441 | | | |
| | | Of-Dxx-511R | GTCTCCACCATCTGTCACAGG | 491 | | | |
| | 4b | Of-dsxb-555P | CAGCAGCTCACCAGTCCTC | 555 | 88 | FAM | BHQ1 |
| | | Of-dsxb-527F | GGAGGCGAGTTCTCAGCTTT | 527 | | | |
| | | Of-dsxb-614R | GGTAGTGCTTGACGTAGGCG | 595 | | | |
| | 5 | Of-dsxb-703P | GCCTCCATACGACCACTACG | 703 | 114 | FAM | BHQ1 |
| | | Of-dsxb-671F | GGTTTCCAGTTCAGTTCCCTACT | 671 | | | |
| | | Of-dsxb-784R | ACAGTCGGAGTAGTAAATGCTGT | 762 | | | |

**Supplemental Table 2.** Primer sequences used for dsRNA synthesis. Lowercase sequence is the T7 RNA polymerase promoter. dsRNA length is given for the target sequence only. RNA molecules also included flanking T7 promoter sequences not included in the length here.

| target | exon | primer name | sequence | position | dsRNA (bp) |
|---|---|---|---|---|---|
| *AmpR* | | T7-AmpR-133F | taatacgactcactatagggATCGAACTGGATCTCAACAG | 133 | 447 |
| | | T7-AmpR-579R | taatacgactcactatagggAGTTAATAGTTTGCGCAACG | 579 | |
| *intersex* | | T7-Of-ix-62F | taatacgactcactatagggTAAGGAAGATGAACATACC | 62 | 161 |
| | | T7-Of-ix-222R | taatacgactcactatagggCACTTTGGATATATTGTCC | 222 | |
| | | T7-Of-ix-f1 | taatacgactcactatagggAGAGTCCCTTGTTGCTACTC | 246 | 201 |
| | | T7-Of-ix-r1 | taatacgactcactatagggGAGCTCTGACTCATGCATTC | 446 | |
| *fruitless* | | T7-Of-fru-181F | taatacgactcactatagggTGGATGTCACTTTGGCTGTC | 181 | 155 |
| | | T7-Of-fru-335R | taatacgactcactatagggAACTGTGACGTCCCTTAGGA | 335 | |
| *dsx* | 1 | T7-Of-dsxc-242F | taatacgactcactatagggTCATGAAGAGAAAGTACTG | 242 | 157 |
| | | T7-Of-dsxc-398R | taatacgactcactatagggCGTTTCTTTACGTTCATCC | 398 | |
| | 2b | T7-Of-dsxc-484F | taatacgactcactataggGTATAGATTGAAGACAAGTC | 484 | 244 |
| | | T7-Of-dsxc-727R | taatacgactcactatagggTAGGATTCCTTATATGTAGG | 727 | |
| | 3b,6 | T7-Of-dsxc-1404F | taatacgactcactataggGAGAACTCTCTTATCAATC | 1404 | 203 |
| | | T7-Of-dsxc-1779R | taatacgactcactatagGGTACAATATATGCTGTAC | 1779 | |
| | 4,5 | T7-Of-dsxc-1448F | taatacgactcactatagggTTAGCTTGCTCAGATTGAC | 1448 | 168 |
| | | T7-Of-dsxc-1615R | taatacgactcactataggGATAAGTTTGGAGGAGTCTC | 1615 | |
| *dmrt99B* | 1 | T7-Of-Dsx-22F | taatacgactcactatagggTGCCGGTGGCGGGACTGCTC | 22 | 88 |
| | | T7-Of-Dsx-109R | taatacgactcactatagggACCTCCTGAGGGCGACCTGG | 109 | |
| | 1 | T7-Of-Dsx-180R | taatacgactcactatagggCGGAAGAAGTGGTGGTTCC | 180 | 159 (w/ 22F) |
| | 2b | T7-Of-Dsx-100F | taatacgactcactatagggCTCAGGAGGTAGCAGGCAC | 100 | 597 |
| | | T7-Of-Dsx-696R | taatacgactcactatagggACTATGGCCTTCCTTCCTTC | 696 | |
| *dmrt93B* | 1 | T7-Of-Dxx-30F | taatacgactcactatagggAGACCGAAGTGCGCCCGCTG | 30 | 56 |
| | | T7-Of-Dxx-85R | taatacgactcactatagggTGCCCTTTCAGCCAGCTGAT | 85 | |
| | 1,2 | T7-Of-Dxx-284R | taatacgactcactatagggCGGTAGAGTGGAAAGAATAG | 284 | 254 (w/ 30F) |

**Supplemental Table 3.** Sample sizes and survival rates for RNA interference experiments. [a] Includes the conserved Med29 domain. [b] Includes the OD1 DNA-binding domain. [c] Includes the OD2 / DMRTA domain.

| target gene | transcript isoforms | region | [dsRNA] (μg/μl) | injected nymphs | surviving adults |
|---|---|---|---|---|---|
| *AmpR* | | 133-579 | 2.5* | 135 | 103 (76.3%) |
| | | | 5.0 | 43 | 40 (93.0%) |
| *ix* | | 62-222 | 1.0 | 68 | 35 (51.5%) |
| | | | 1.5 | 35 | 26 (74.3%) |
| | | | 2.5 | 30 | 11 (36.7%) |
| | | 246-446 [a] | 1.0 | 102 | 44 (43.1%) |
| | | | 1.5 | 40 | 0 |
| | | | 2.5 | 30 | 0 |
| *fru* | | 181-335 | 1.5 | 28 | 22 (78.6%) |
| | | | 2.5 | 39 | 21 (53.8%) |
| *dsx* | all | 242-398 [b] | 2.5 | 157 | 122 (77.7%) |
| | | | 3.0 | 36 | 27 (75.0%) |
| | *E* | 484-727 | 2.5 | 113 | 83 (73.5%) |
| | *D* | 1404-1779 | 2.5 | 86 | 62 (72.1%) |
| | *A, B, C* | 1448-1615 [c] | 2.5 | 92 | 58 (63.0%) |
| *dmrt99B* | all | 22-109 [b] | 2.5 | 10 | 10 (100%) |
| | | | 3.0 | 38 | 24 (63.2%) |
| | | | 3.5 | 33 | 19 (57.6%) |
| | all | 22-180 [b] | 2.5 | 44 | 39 (88.6%) |
| | *C* | 100-696 | 2.5 | 49 | 42 (85.7%) |
| *dmrt93B* | all | 30-85 [b] | 3.0 | 45 | 31 (68.9%) |
| | | | 3.5 | 32 | 15 (46.9%) |
| | all | 30-284 | 2.5 | 44 | 35 (79.5%) |
| *dsx, dmrt99B, dmrt93B* co-injected dsRNAs | | [b] | 3.0 (1.0 each) | 10 | 8 (80.0%) |
| | | | 3.5 (1.17 each) | 20 | 20 (100%) |
| | | | 7.2 (2.4 each) | 37 | 19 (51.4%) |
| *dsx, dmrt99B, dmrt93B* tandem dsRNA | | [b] | 1.0 | 20 | 17 (85.0%) |
| | | | 2.5 | 20 | 17 (85.0%) |
| | | | 3.0 | 18 | 9 (50.0%) |
| | | | 5.0 | 20 | 10 (50.0%) |
| | | | 6.0 | 48 | 24 (50.0%) |

### Supplemental Figure Legends

**Supplemental Figure S1**

**Consensus phylogram for DMRT proteins**. Amino acid sequences were aligned by Clustal-Omega and maximum likelihood trees were generated by RAxML with bootstrap resampling. Tips are labeled with the species name and GenBank accession number of each sequence. Bootstrap support is indicated distal of each node. The phylogram is rooted with DMRT family proteins from non-insect species. Sequences from *Oncopeltus fasciatus* are highlighted in orange. Sequences from *Drosophila melanogaster* are highlighted in green. The four presumptive paralog groups are bracketed by heavy vertical bars at the right.

**Supplemental Figure S2**

**Consensus phylogram for Doublesex proteins**. The phylogram is rooted with the Dsx sequence from *Daphnia magna*.

**Supplemental Figure S3**

**Consensus phylogram for Fruitless proteins**. The phylogram is rooted with homologs from Collembola.

**Supplemental Figure S4**

**Consensus phylogram for Intersex proteins**. The phylogram is rooted with homologs from Chelicerata.

**Supplemental Figure S5**

**Validation of RNA interference.** Gene expression was measured by qRT-PCR in specimens treated with a control dsRNA sequence encoding *Ampicillin Resistance* (*AmpR*) and with dsRNA targeting genes of interest. Female and male specimens were validated separately. Sample sizes for each group are indicated at the top of each panel. Dots indicate individual measurements and are summarized by Tukey's box-whisker plots showing median (heavy bar), interquartile range (box) and full range (whiskers). Significant difference from the control measurements is indicated by the *p*-value given in red, which comes from a Wilcoxon rank sum test with FDR correction.

**Supplemental Figure S6**

**Structure and expression of *dmrt99B* and *dmrt93B* in *O. fasciatus*.** (a) Orthologs were identified on different genomic scaffolds in the *O. fasciatus* genome. Boxes represent exons; angled lines represent introns. Introns were very long compared to neighboring gene predictions. Flags indicate the start of transcription. (b) Transcripts from *dmrt99B* and *dmrt93B*. The first exon of each transcript is aligned at the left. Omitted exons are left empty. Alternative start or stop sites for exons are indicated by lowercase letters (e.g. 2a, 2b) Lighter shading indicates UTRs. (c) A heatmap shows the relative transcript abundance from five body regions in each sex for *dmrt93B* and *dmrt99B*. *dmrt99B* was not detected in any male tissues, but showed relatively consistent, low expression in female tissues, excluding the ovipositor. We failed to detect *dmrt99B* exon 2, suggesting that all detectable expression in the sampled tissues reflects expression of transcript isoform B. Expression of *dmrt93B* was high in the gonads of both sexes, but low or undetectable in other tissues. This expression was greater in testes than in ovaries. We did not detect any expression of *dmrt93B* exon 4a in females, suggesting male specificity in transcript isoform B. Each panel represents the mean of 5 to 7 independent measurements. Expression differences between females and males were tested using the Wilcoxon rank sum test (* indicates $p < 0.05$).

**Supplemental Figure S7**

**Gonad anatomy in RNAi specimens.** To examine potential RNAi effects on gonad anatomy, adults were dissected by removing the ventral cuticle. In each panel, anterior is up, and the scale is in mm units. The gonad on the left is outlined. Females (top row) have two ovaries, which each consist of seven ovarioles that lead to an oviduct. Males (bottom row) have two testes consisting of seven testioles, connecting to a vas deferens. No obvious anatomical differences are apparent in unmanipulated or dsRNA-treated specimens.

**Supplemental Figure S8**

***intersex* promotes *fruitless* expression in males.** Transcription-level gene interactions were tested by measuring expression using qRT-PCR in an RNAi background. *fruitless* expression was measured in control males (*AmpR* dsRNA), in *ix* RNAi males and in *fru* RNAi males. *fru* expression was reduced relative to controls in *fru* and *ix* knockdown, suggesting that *ix* promotes *fru* expression in the wildtype. Dots indicate individual measurements and are summarized by Tukey's box-whisker plots. Significant difference from the mean of control measurements is indicated by the *p*-value given in red, which comes from a Wilcoxon rank sum test with FDR correction.

**Supplemental Figure S9**

**Relationships between genitalia length and fourth sternite curvature.** Dots represent individual specimens, colored by presumptive sex. While each sex has a different average value for each trait, there is little correlation between traits within each group.

**Supplemental Figure S10**

**Relationships fourth sternite curvature and abdominal melanism.** Dots represent individual specimens, colored by presumptive sex. While each sex has a different average value for each trait, there is little correlation between traits within each group.

**Supplemental Figure S11**

**Quantification of RNAi effects, including *dmrt99B* and *dmrt93B***, on genitalia length (a-b), sternite curvature (d-e) and abdominal melanism (g-h) in females (a,d,g) and males (b,e,h). Dots represent individual specimens. Gray outlines show the relative distribution of values. The mean of control (*AmpR* dsRNA) specimens is indicated by the horizontal line in each panel. Stars indicate significant difference from control values, based on one-sided Wilcoxon rank sum tests with FDR correction. * $p < 0.05$; ** $p < 0.01$; *** $p < 0.001$, and are summarized beside a sketch of typically female and male phenotypes at the right (c,f,i).

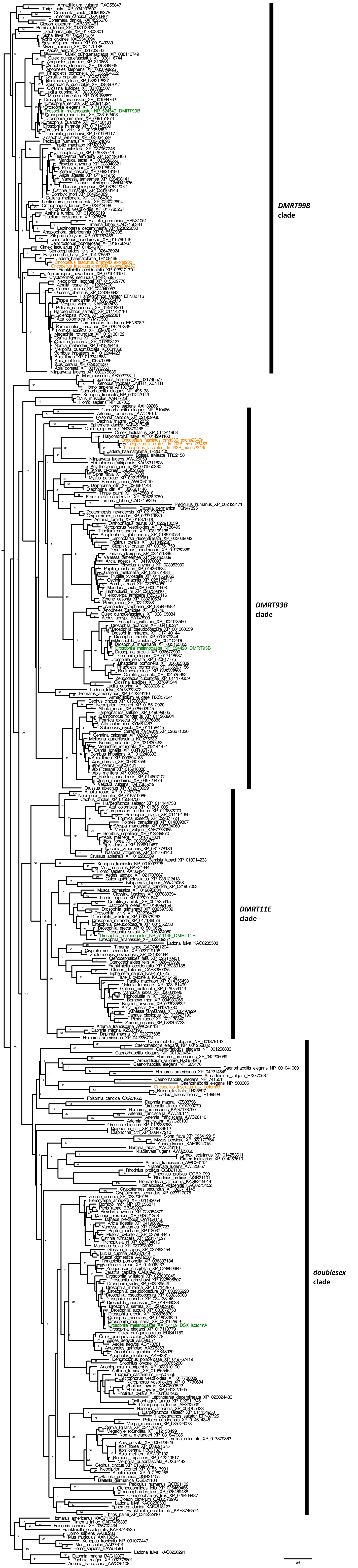
DMRT99B
clade
DMRT93B
clade
DMRT11E
clade
doublesex
clade
0.6

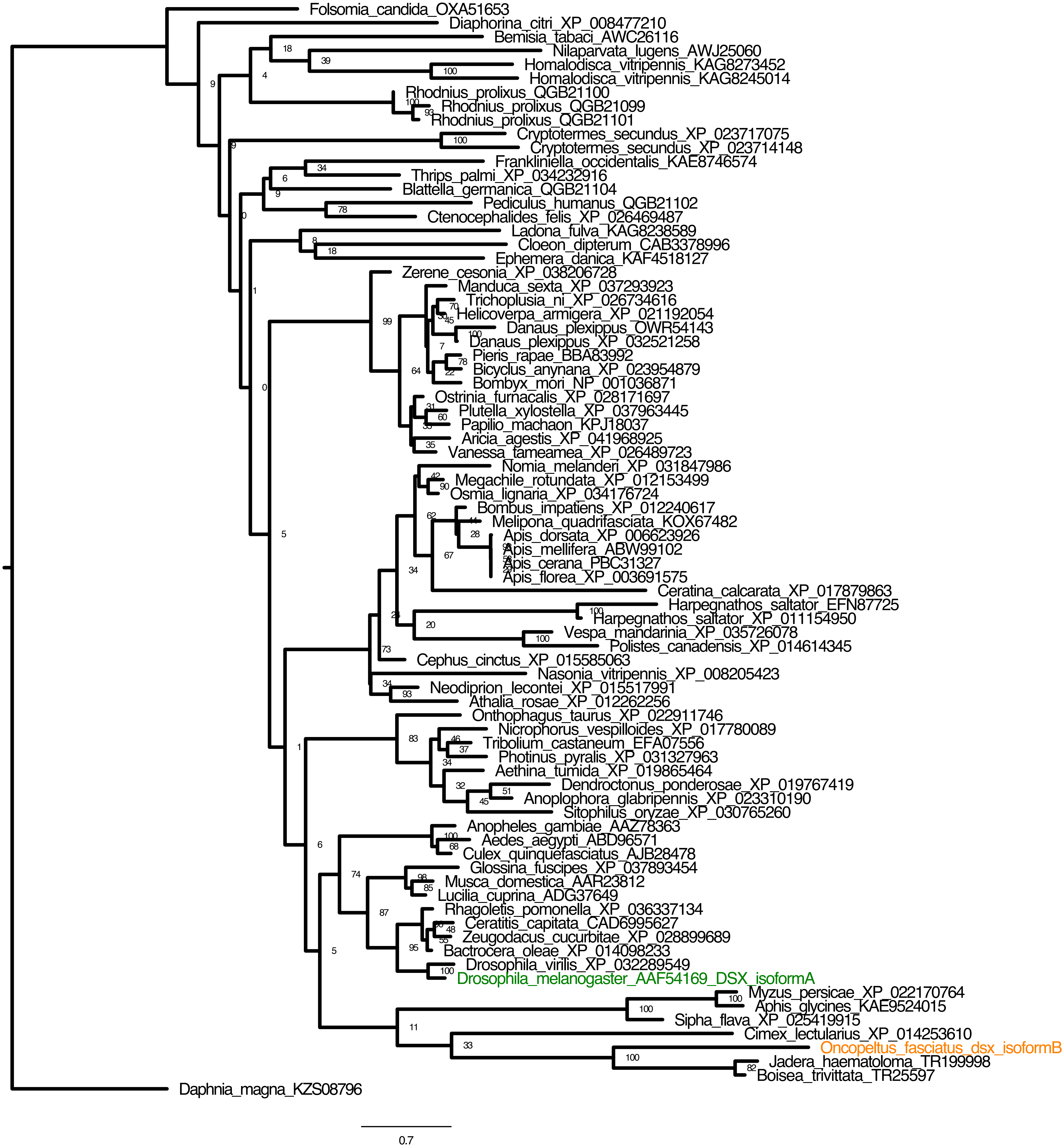

Folsomia_candida_OXA51653
Diaphorina_citri_XP_008477210
Bemisia_tabaci_AWC26116
Nilaparvata_lugens_AWJ25060
Homalodisca_vitripennis_KAG8273452
Homalodisca_vitripennis_KAG8245014
Rhodnius_prolixus_QGB21100
Rhodnius_prolixus_QGB21099
Rhodnius_prolixus_QGB21101
Cryptotermes_secundus_XP_023717075
Cryptotermes_secundus_XP_023714148
Frankliniella_occidentalis_KAE8746574
Thrips_palmi_XP_034232916
Blattella_germanica_QGB21104
Pediculus_humanus_QGB21102
Ctenocephalides_felis_XP_026469487
Ladona_fulva_KAG8238589
Cloeon_dipterum_CAB3378996
Ephemera_danica_KAF4518127
Zerene_cesonia_XP_038206728
Manduca_sexta_XP_037293923
Trichoplusia_ni_XP_026734616
Helicoverpa_armigera_XP_021192054
Danaus_plexippus_OWR54143
Danaus_plexippus_XP_032521258
Pieris_rapae_BBA83992
Bicyclus_anynana_XP_023954879
Bombyx_mori_NP_001036871
Ostrinia_furnacalis_XP_028171697
Plutella_xylostella_XP_037963445
Papilio_machaon_KPJ18037
Aricia_agestis_XP_041968925
Vanessa_tameamea_XP_026489723
Nomia_melanderi_XP_031847986
Megachile_rotundata_XP_012153499
Osmia_lignaria_XP_034176724
Bombus_impatiens_XP_012240617
Melipona_quadrifasciata_KOX67482
Apis_dorsata_XP_006623926
Apis_mellifera_ABW99102
Apis_cerana_PBC31327
Apis_florea_XP_003691575
Ceratina_calcarata_XP_017879863
Harpegnathos_saltator_EFN87725
Harpegnathos_saltator_XP_011154950
Vespa_mandarinia_XP_035726078
Polistes_canadensis_XP_014614345
Cephus_cinctus_XP_015585063
Nasonia_vitripennis_XP_008205423
Neodiprion_lecontei_XP_015517991
Athalia_rosae_XP_012262256
Onthophagus_taurus_XP_022911746
Nicrophorus_vespilloides_XP_017780089
Tribolium_castaneum_EFA07556
Photinus_pyralis_XP_031327963
Aethina_tumida_XP_019865464
Dendroctonus_ponderosae_XP_019767419
Anoplophora_glabripennis_XP_023310190
Sitophilus_oryzae_XP_030765260
Anopheles_gambiae_AAZ78363
Aedes_aegypti_ABD96571
Culex_quinquefasciatus_AJB28478
Glossina_fuscipes_XP_037893454
Musca_domestica_AAR23812
Lucilia_cuprina_ADG37649
Rhagoletis_pomonella_XP_036337134
Ceratitis_capitata_CAD6995627
Zeugodacus_cucurbitae_XP_028899689
Bactrocera_oleae_XP_014098233
Drosophila_virilis_XP_032289549
Drosophila_melanogaster_AAF54169_DSX_isoformA
Myzus_persicae_XP_022170764
Aphis_glycines_KAE9524015
Sipha_flava_XP_025419915
Cimex_lectularius_XP_014253610
Oncopeltus_fasciatus_dsx_isoformB
Jadera_haematoloma_TR199998
Boisea_trivittata_TR25597
Daphnia_magna_KZS08796
0.7

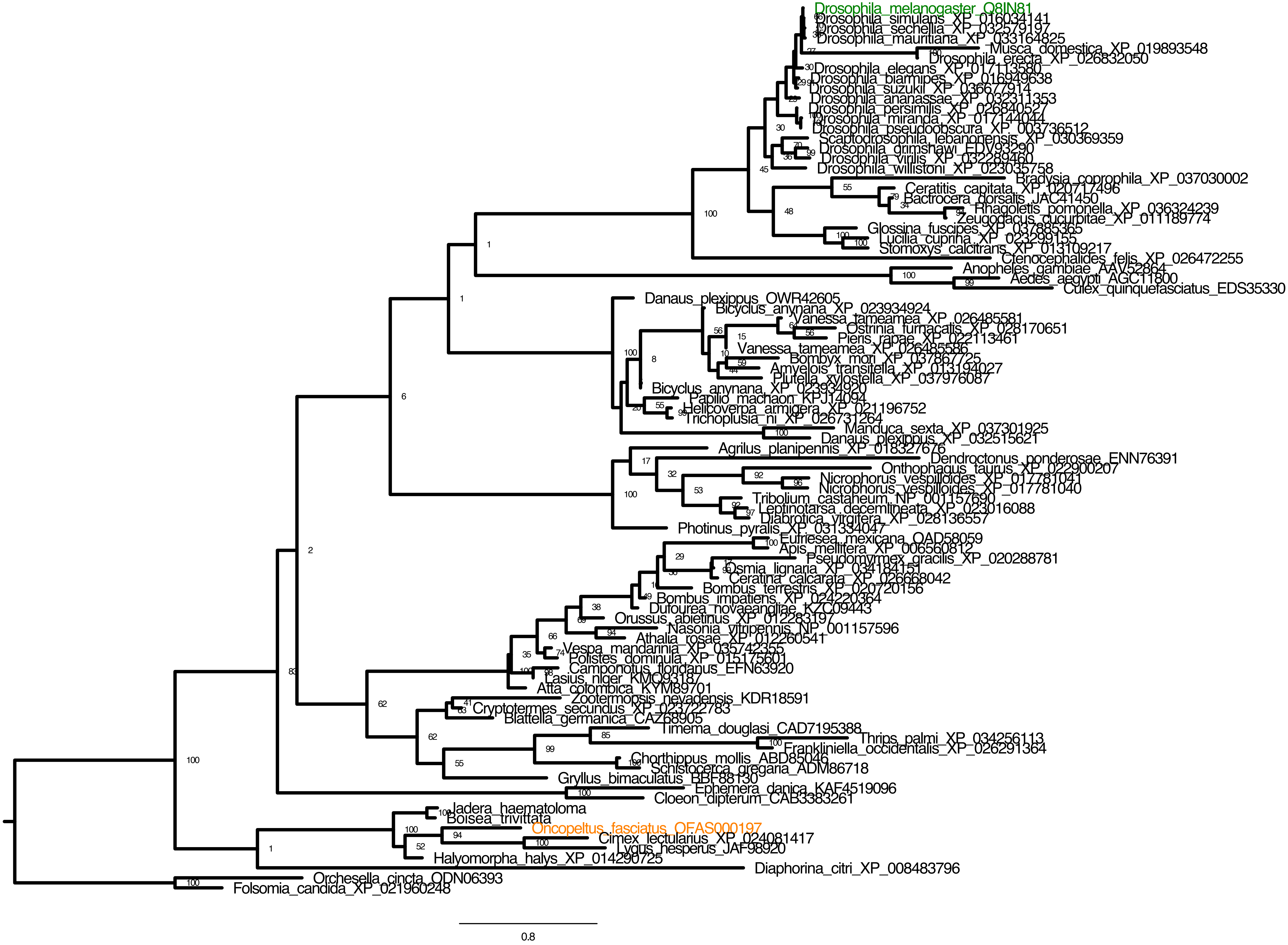

Drosophila_melanogaster_Q8IN81
Drosophila_simulans_XP_016034141
Drosophila_sechellia_XP_032579197
Drosophila_mauritiana_XP_033164825
Musca_domestica_XP_019893548
Drosophila_erecta_XP_026832050
Drosophila_elegans_XP_017113580
Drosophila_biarmipes_XP_016949638
Drosophila_suzukii_XP_036677914
Drosophila_ananassae_XP_032311353
Drosophila_persimilis_XP_026840527
Drosophila_miranda_XP_017144044
Drosophila_pseudoobscura_XP_003736512
Scaptodrosophila_lebanonensis_XP_030369359
Drosophila_grimshawi_EDV93290
Drosophila_virilis_XP_032289460
Drosophila_willistoni_XP_023035758
Bradysia_coprophila_XP_037030002
Ceratitis_capitata_XP_020717496
Bactrocera_dorsalis_JAC41450
Rhagoletis_pomonella_XP_036324239
Zeugodacus_cucurbitae_XP_011189774
Glossina_fuscipes_XP_037885365
Lucilia_cuprina_XP_023299155
Stomoxys_calcitrans_XP_013109217
Ctenocephalides_felis_XP_026472255
Anopheles_gambiae_AAV52864
Aedes_aegypti_AGC11800
Culex_quinquefasciatus_EDS35330
Danaus_plexippus_OWR42605
Bicyclus_anynana_XP_023934924
Vanessa_tameamea_XP_026485581
Ostrinia_furnacalis_XP_028170651
Pieris_rapae_XP_022113461
Vanessa_tameamea_XP_026485586
Bombyx_mori_XP_037867725
Amyelois_transitella_XP_013194027
Plutella_xylostella_XP_037976087
Bicyclus_anynana_XP_023934920
Papilio_machaon_KPJ14094
Helicoverpa_armigera_XP_021196752
Trichoplusia_ni_XP_026731264
Manduca_sexta_XP_037301925
Danaus_plexippus_XP_032515621
Agrilus_planipennis_XP_018327676
Dendroctonus_ponderosae_ENN76391
Onthophagus_taurus_XP_022900207
Nicrophorus_vespilloides_XP_017781041
Nicrophorus_vespilloides_XP_017781040
Tribolium_castaneum_NP_001157690
Leptinotarsa_decemlineata_XP_023016088
Diabrotica_virgifera_XP_028136557
Photinus_pyralis_XP_031334047
Eufriesea_mexicana_OAD58059
Apis_mellifera_XP_006560812
Pseudomyrmex_gracilis_XP_020288781
Osmia_lignaria_XP_034184151
Ceratina_calcarata_XP_026668042
Bombus_terrestris_XP_020720156
Bombus_impatiens_XP_024220364
Dufourea_novaeangliae_KZC09443
Orussus_abietinus_XP_012283197
Nasonia_vitripennis_NP_001157596
Athalia_rosae_XP_012260541
Vespa_mandarinia_XP_035742355
Polistes_dominula_XP_015175601
Camponotus_floridanus_EFN63920
Lasius_niger_KMQ93187
Atta_colombica_KYM89701
Zootermopsis_nevadensis_KDR18591
Cryptotermes_secundus_XP_023722783
Blattella_germanica_CAZ68905
Timema_douglasi_CAD7195388
Thrips_palmi_XP_034256113
Frankliniella_occidentalis_XP_026291364
Chorthippus_mollis_ABD85046
Schistocerca_gregaria_ADM86718
Gryllus_bimaculatus_BBF88130
Ephemera_danica_KAF4519096
Cloeon_dipterum_CAB3383261
Jadera_haematoloma
Boisea_trivittata
Oncopeltus_fasciatus_OFAS000197
Cimex_lectularius_XP_024081417
Lygus_hesperus_JAF98920
Halyomorpha_halys_XP_014290725
Diaphorina_citri_XP_008483796
Orchesella_cincta_ODN06393
Folsomia_candida_XP_021960248
0.8

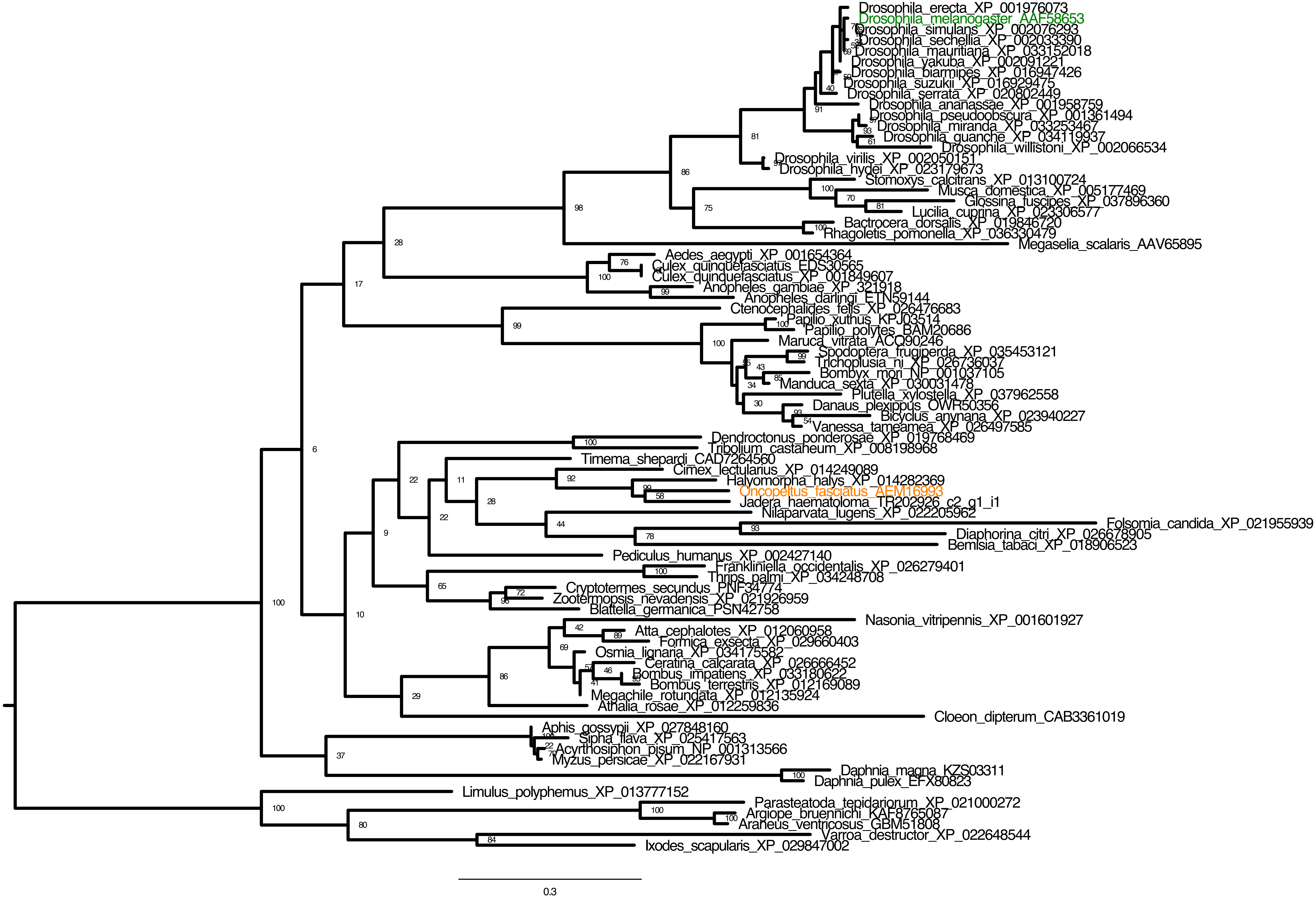

Drosophila_erecta_XP_001976073
Drosophila_melanogaster_AAF58653
Drosophila_simulans_XP_002076293
Drosophila_sechellia_XP_002033390
Drosophila_mauritiana_XP_033152018
Drosophila_yakuba_XP_002091221
Drosophila_biarmipes_XP_016947426
Drosophila_suzukii_XP_016929475
Drosophila_serrata_XP_020802449
Drosophila_ananassae_XP_001958759
Drosophila_pseudoobscura_XP_001361494
Drosophila_miranda_XP_033253467
Drosophila_guanche_XP_034119937
Drosophila_willistoni_XP_002066534
Drosophila_virilis_XP_002050151
Drosophila_hydei_XP_023179673
Stomoxys_calcitrans_XP_013100724
Musca_domestica_XP_005177469
Glossina_fuscipes_XP_037896360
Lucilia_cuprina_XP_023306577
Bactrocera_dorsalis_XP_019846720
Rhagoletis_pomonella_XP_036330479
Megaselia_scalaris_AAV65895
Aedes_aegypti_XP_001654364
Culex_quinquefasciatus_EDS30565
Culex_quinquefasciatus_XP_001849607
Anopheles_gambiae_XP_321918
Anopheles_darlingi_ETN59144
Ctenocephalides_felis_XP_026476683
Papilio_xuthus_KPJ03514
Papilio_polytes_BAM20686
Maruca_vitrata_ACQ90246
Spodoptera_frugiperda_XP_035453121
Trichoplusia_ni_XP_026736037
Bombyx_mori_NP_001037105
Manduca_sexta_XP_030031478
Plutella_xylostella_XP_037962558
Danaus_plexippus_OWR50356
Bicyclus_anynana_XP_023940227
Vanessa_tameamea_XP_026497585
Dendroctonus_ponderosae_XP_019768469
Tribolium_castaneum_XP_008198968
Timema_shepardi_CAD7264560
Cimex_lectularius_XP_014249089
Halyomorpha_halys_XP_014282369
Oncopeltus_fasciatus_AEM16993
Jadera_haematoloma_TR202926_c2_g1_i1
Nilaparvata_lugens_XP_022205962
Folsomia_candida_XP_021955939
Diaphorina_citri_XP_026678905
Bemisia_tabaci_XP_018906523
Pediculus_humanus_XP_002427140
Frankliniella_occidentalis_XP_026279401
Thrips_palmi_XP_034248708
Cryptotermes_secundus_PNF34774
Zootermopsis_nevadensis_XP_021926959
Blattella_germanica_PSN42758
Nasonia_vitripennis_XP_001601927
Atta_cephalotes_XP_012060958
Formica_exsecta_XP_029660403
Osmia_lignaria_XP_034175582
Ceratina_calcarata_XP_026666452
Bombus_impatiens_XP_033180622
Bombus_terrestris_XP_012169089
Megachile_rotundata_XP_012135924
Athalia_rosae_XP_012259836
Cloeon_dipterum_CAB3361019
Aphis_gossypii_XP_027848160
Sipha_flava_XP_025417563
Acyrthosiphon_pisum_NP_001313566
Myzus_persicae_XP_022167931
Daphnia_magna_KZS03311
Daphnia_pulex_EFX80823
Limulus_polyphemus_XP_013777152
Parasteatoda_tepidariorum_XP_021000272
Argiope_bruennichi_KAF8765087
Araneus_ventricosus_GBM51808
Varroa_destructor_XP_022648544
Ixodes_scapularis_XP_029847002
0.3

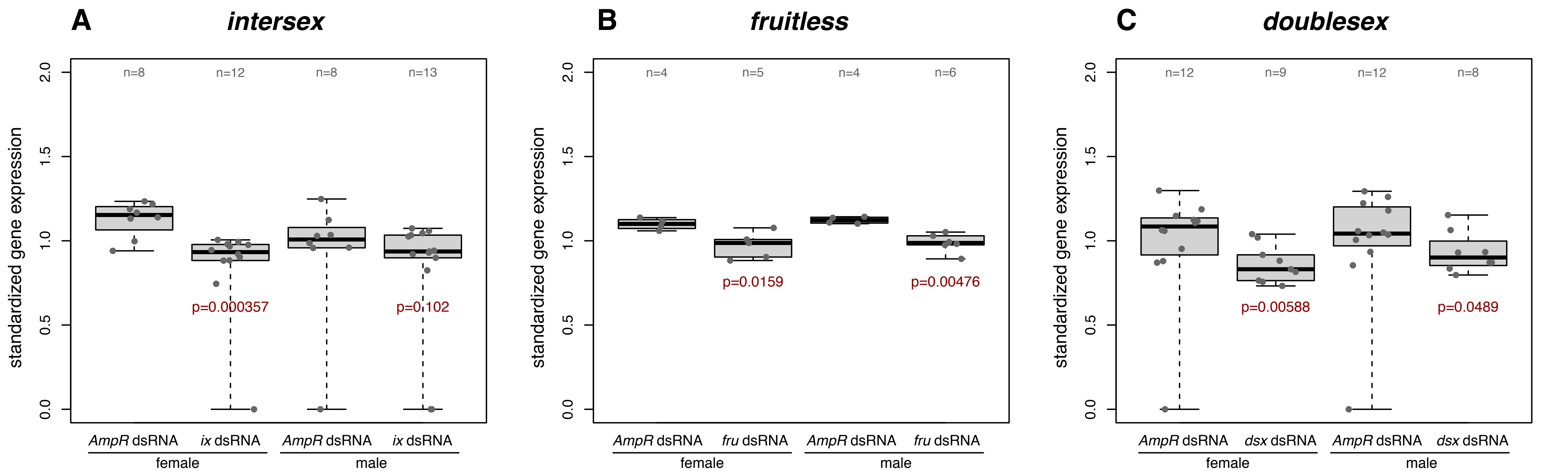
A
intersex
B
fruitless
C
doublesex
standardized gene expression
0.0
0.5
1.0
1.5
2.0
n=8
n=12
n=8
n=13
p=0.000357
p=0.102
n=4
n=5
n=4
n=6
p=0.0159
p=0.00476
n=12
n=9
n=12
n=8
p=0.00588
p=0.0489
AmpR dsRNA
ix dsRNA
fru dsRNA
dsx dsRNA
female
male

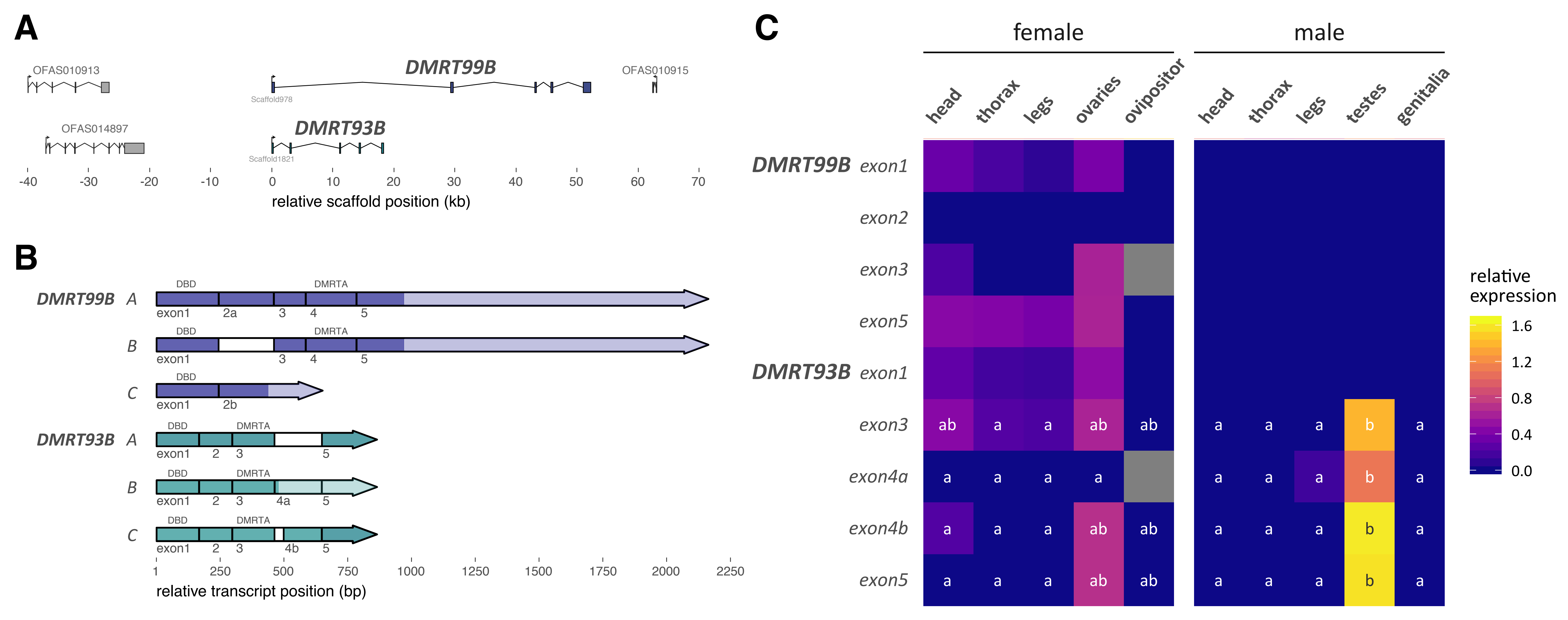
A
OFAS010913
DMRT99B
OFAS010915
Scaffold978
OFAS014897
DMRT93B
Scaffold1821
-40 -30 -20 -10 0 10 20 30 40 50 60 70
relative scaffold position (kb)
B
DMRT99B A
DBD
DMRTA
exon1 2a 3 4 5
B
exon1 3 4 5
C
exon1 2b
DMRT93B A
exon1 2 3 5
B
exon1 2 3 4a 5
C
exon1 2 3 4b 5
1 250 500 750 1000 1250 1500 1750 2000 2250
relative transcript position (bp)
C
female
male
head thorax legs ovaries ovipositor
head thorax legs testes genitalia
DMRT99B exon1
exon2
exon3
exon5
DMRT93B exon1
exon3
exon4a
exon4b
exon5
relative expression
1.6
1.2
0.8
0.4
0.0

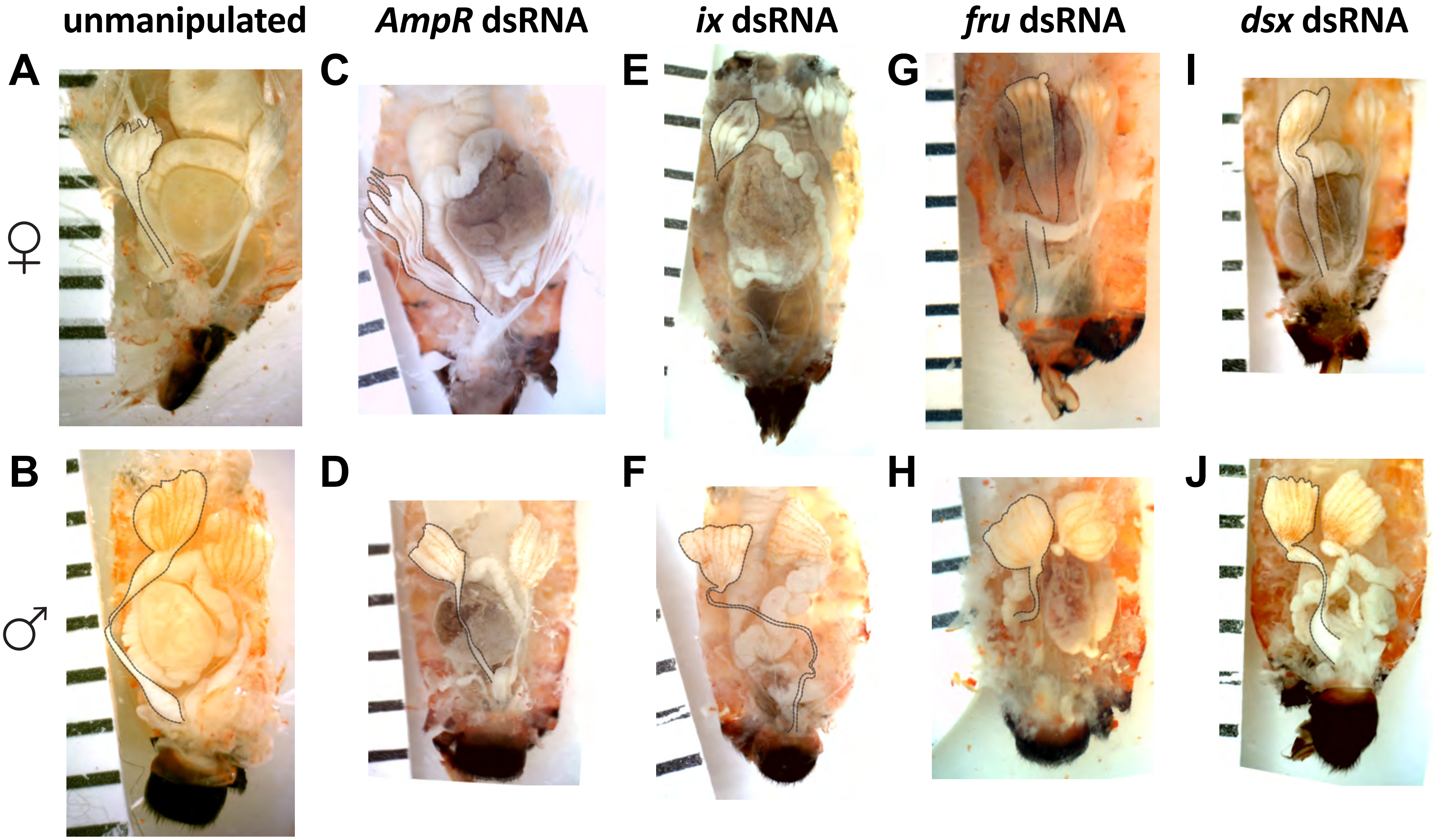
unmanipulated
AmpR dsRNA
ix dsRNA
fru dsRNA
dsx dsRNA
A
B
C
D
E
F
G
H
I
J
♀
♂

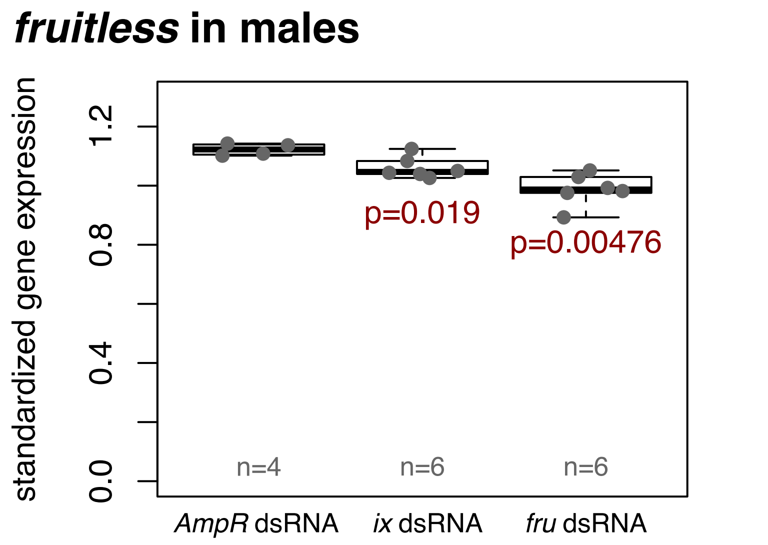

*fruitless* in males
standardized gene expression
1.2
0.8
0.4
0.0
p=0.019
p=0.00476
n=4
n=6
n=6
*AmpR* dsRNA
*ix* dsRNA
*fru* dsRNA

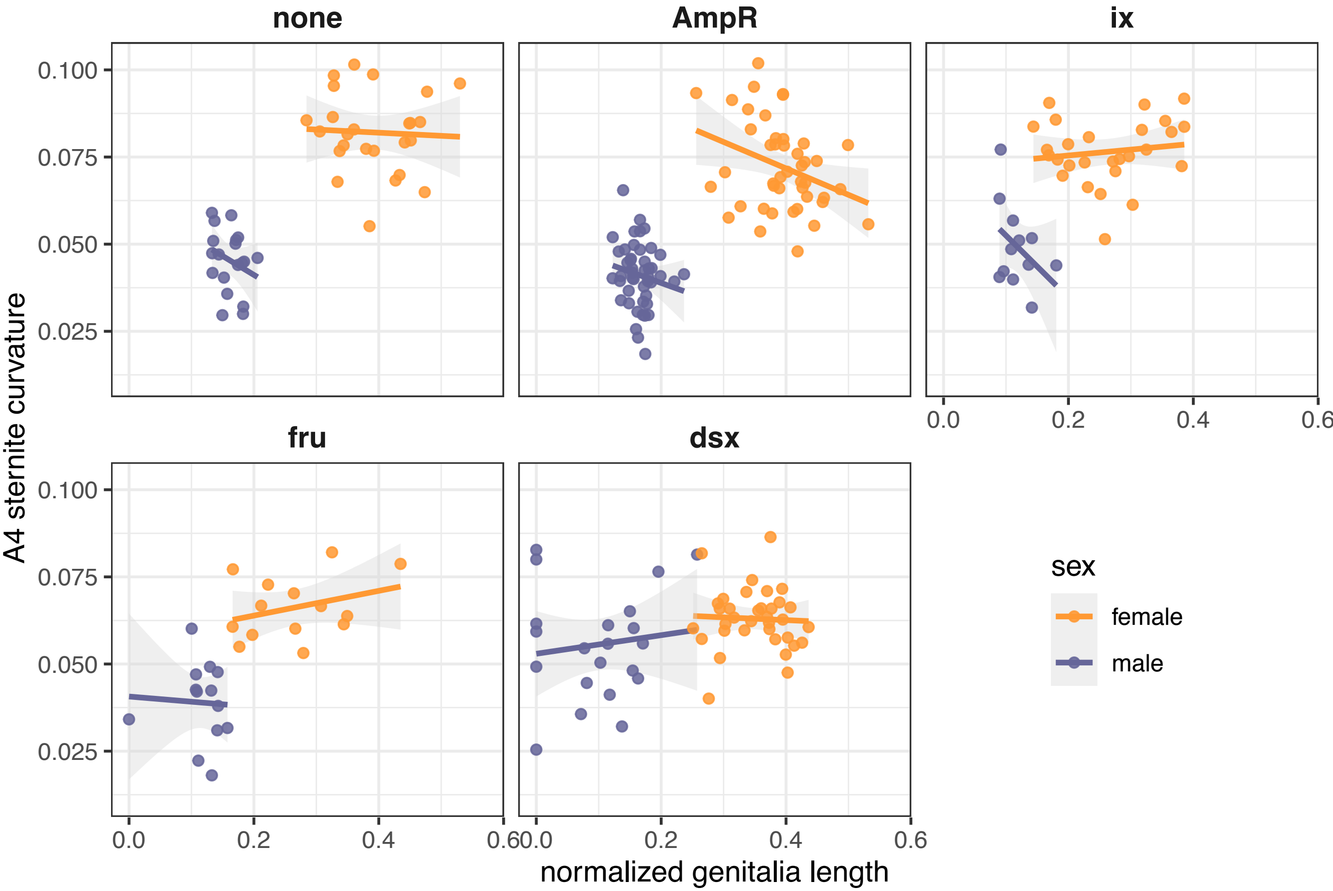

none
AmpR
ix
fru
dsx
A4 sternite curvature
normalized genitalia length
0.100
0.075
0.050
0.025
0.0
0.2
0.4
0.6
sex
female
male

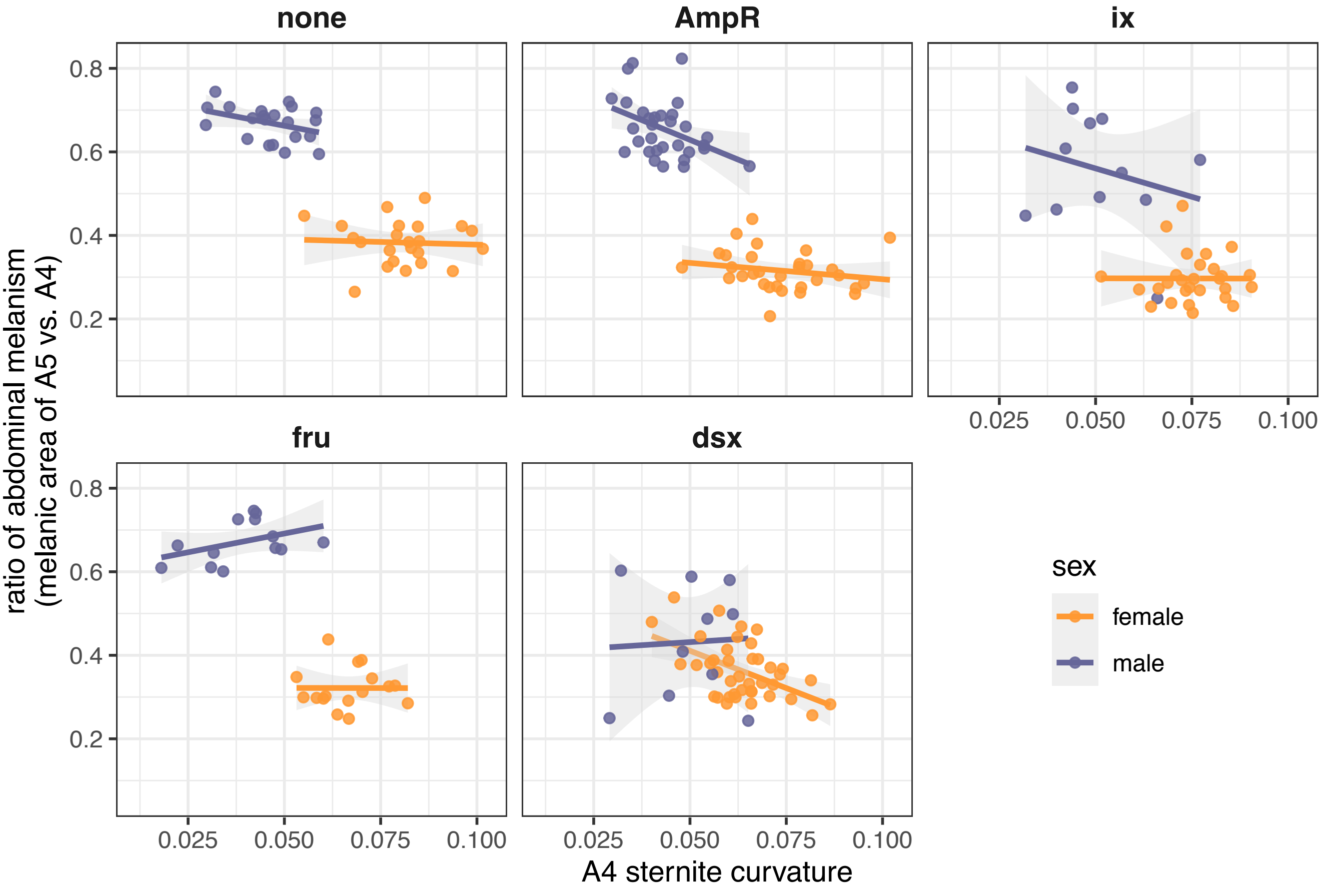

none
AmpR
ix
fru
dsx
0.8
0.6
0.4
0.2
0.025
0.050
0.075
0.100
ratio of abdominal melanism
(melanic area of A5 vs. A4)
A4 sternite curvature
sex
female
male

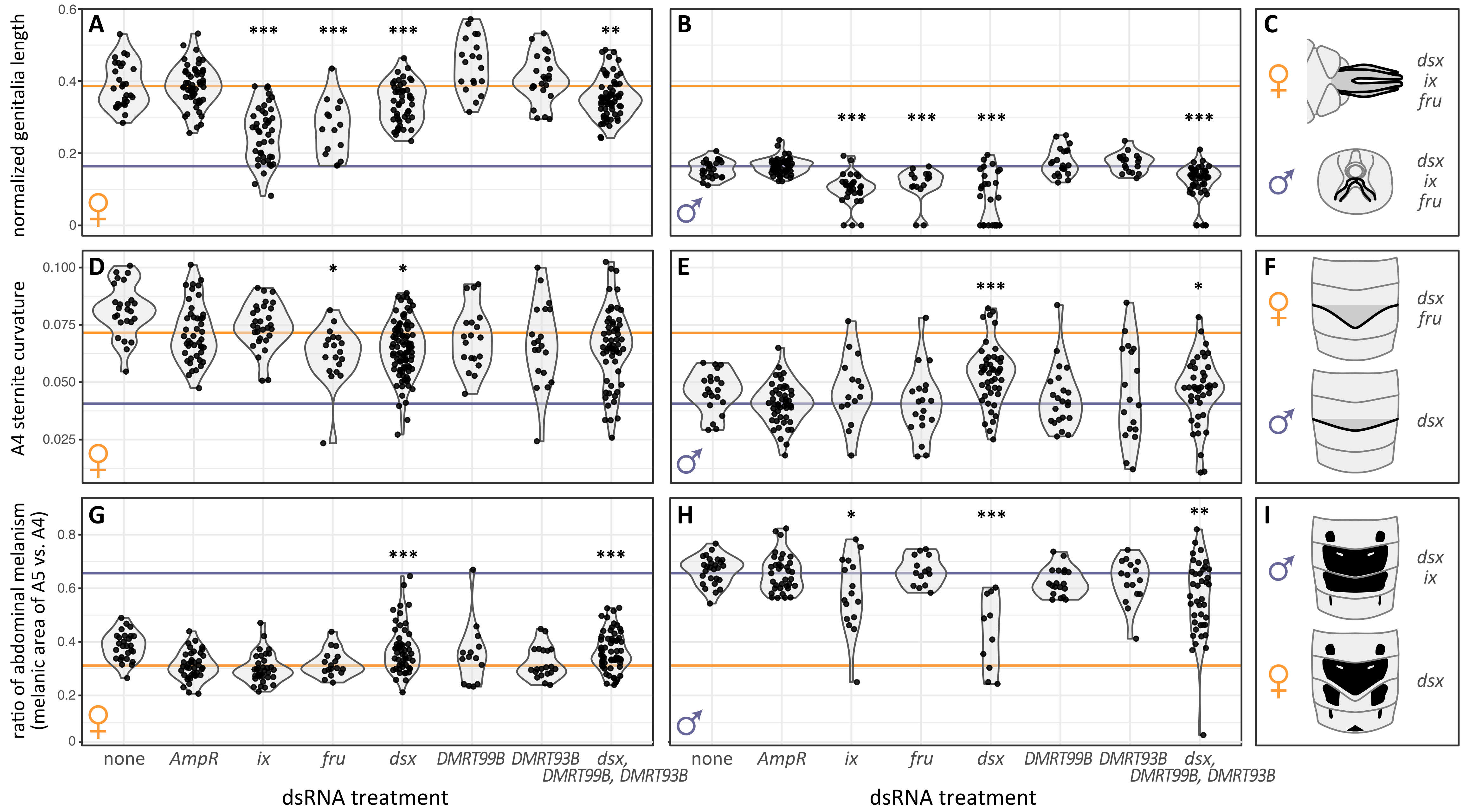
A
B
C
D
E
F
G
H
I
normalized genitalia length
A4 sternite curvature
ratio of abdominal melanism
(melanic area of A5 vs. A4)
***
**
*
dsx
ix
fru
0.6
0.4
0.2
0
0.100
0.075
0.050
0.025
0.8
none
AmpR
DMRT99B
DMRT93B
dsx,
DMRT99B, DMRT93B
dsRNA treatment
